# Resistance Signatures Manifested in Early Drug Response in Cancer and Across Species

**DOI:** 10.1101/2024.07.05.602281

**Authors:** Cole Ruoff, Allision Mitchell, Priya Mondal, Vishaka Gopalan, Arashdeep Singh, Michael Gottesman, Sridhar Hannenhalli

**Affiliations:** Cancer Data Science Lab, Center for Cancer Research, National Cancer Institute, National Institutes of Health, Bethesda, Maryland, 20892, USA; Laboratory of Cell Biology, Center for Cancer Research, National Cancer Institute, National Institutes of Health, Bethesda, Maryland, 20892, USA

## Abstract

Therapeutic resistance is a major cause of cancer treatment failure, with increasing evidence suggesting a non-genetic basis. This non-genetic resistance is often due to drug-resistant transcriptional cell states, either induced by treatment or pre-existing in some cells. However, the connection between early cellular drug response and long-term resistance is poorly understood. Moreover, it is unknown whether resistance-associated early transcriptional responses are evolutionarily conserved. Integrating long-term drug resistance and early drug response data across multiple cancer cell lines, bacteria, and yeast, our findings indicate that cancer states in drug-naive populations and shortly after treatment share transcriptional properties with fully resistant populations, some of which are evolutionarily conserved. CRISPR-Cas9 knockout of resistant states’ markers increased sensitivity to Prexasertib in ovarian cancer cells. Finally, early resistant state signatures discriminated therapy responders from non-responders across multiple human cancer trials, and distinguished premalignant breast lesions that progress to malignancy from those that do not.

## INTRODUCTION

While numerous chemotherapeutic, targeted, and monoclonal antibody-based drugs are approved for various cancers, resistance to these drugs invariably emerges resulting in relapse and eventual death of the patient. Indeed, therapeutic resistance is the leading cause of treatment failure and cancer mortality^1^. Several mechanisms driving resistance have been identified^2^. While mutation-driven resistance has been the predominant paradigm thus far^3^, more recent evidence points to an epigenetic, or transcriptional basis of resistance, whereby resistance results from a subpopulation of malignant cells occupying drug resistant transcriptional states that allow for their survival through the treatment^4,5^. Pre-existing, reversible resistant cells are referred to as drug tolerant persister (DTP) cells^6^ distinguishing them from the emergent, stable, resistant states^7^.

Recent literature has identified and characterized these resistant cell states^8,9^ but much is unknown about the timeline in which these states develop, which common transcriptional aspects of long-term resistance are simply non-specific cellular responses to any drug treatment, and whether these aspects are shared with our unicellular ancestors. For instance, it is not entirely clear whether and to what extent long-term resistance is evident soon after drug treatment. Furthermore, these immediate drug responses leading up to long-term drug resistance may be an inherent property of cells shared with unicellular ancestors^10^. Here, we investigate whether (i) there is a similarity between early transcriptional response to drugs and long-term resistance, (ii) these responses are evolutionarily conserved, and (iii) resistance-associated early transcriptional response has clinical implications.

Toward these aims, we first derived a long-term broad resistance signature from published studies of resistance across multiple cell lines and drugs^9^. We also obtained a large scRNA-seq dataset consisting of 581,308 cells in three well characterized cancer cell lines (A549, K562, MCF7) treated with 188 different compounds at 24 hours post-treatment^11^. These data represent the early transcriptional response to drugs. We then compared early response single-cell data with the long-term resistance-associated transcriptional signature to reveal aspects of long-term resistance manifested in early response. We additionally assessed the evolutionary conservation of resistance-associated early response to drugs based on previously published resistance-associated transcriptional response datasets in the bacterium *Escherichia coli*^12^ and the fungus *Candida auris*^13^.

Our findings suggest that transcriptional states associated with long-term resistance are manifested early on after the drug treatment; across all three cell lines investigated here, we observed early response transcriptional states similar to long-term resistant states. Furthermore, our analyses suggest that these early harbingers of long-term resistance may partly exist prior to drug treatment as well as be induced by drug treatment. These pre-existing and early drug response states share characteristics across the cell lines, including high oxidative phosphorylation, EMT, hypoxia, and MYC activity. Interestingly, similar states emerge in *E. coli* and *C. auris*, after multiple drug treatments, reflecting that this mode of resistance is an evolutionarily conserved mechanism. Finally, we find that the resistance-associated early transcriptional response signatures have prognostic value in multiple clinical contexts and in multiple cancer types, in terms of discriminating responders from non-responders, as well as discriminating premalignant breast lesions that progress to malignancy from those that do not.

## RESULTS

### Multiple cell states in pre-treatment and early response data express long-term resistance programs

To identify and characterize early post-treatment manifestation of long-term drug resistance, we first set out to create a robust gene signature of long-term resistance. We aimed to derive this signature from multiple cancer types and drug treatments to characterize common broadly applicable resistance signatures and mechanisms. To this end, we compiled a drug resistance gene signature composed of common upregulated genes (Table S1) in six different drug-resistant samples drawn from previously published studies devised specifically to characterize cancer samples with fully established drug resistance^9^ (Methods). These long-term resistance experiments were done in breast cancer, melanoma, and colorectal cancer cell lines, and thus the derived gene set represents a general drug resistance program. Supporting the broader applicability of the resistance signature, we further validated the gene signature in an additional dataset comprising multiple timepoints throughout resistance development in yet another cancer type –– lung adenocarcinoma^9^. Using gene set enrichment analysis, we calculated the enrichment of the resistance signature among the upregulated genes at each time point (Methods). Encouragingly, we saw a strong enrichment at day 14 (Figure 1A), thus supporting the broader validity of the resistance signature. Functional enrichment analysis of the resistance signature (Figure S1A) revealed enrichment of known resistance associated pathways such as EMT and hypoxia^14,15^ further supporting the signature’s role in resistance.

**Figure 1A:**
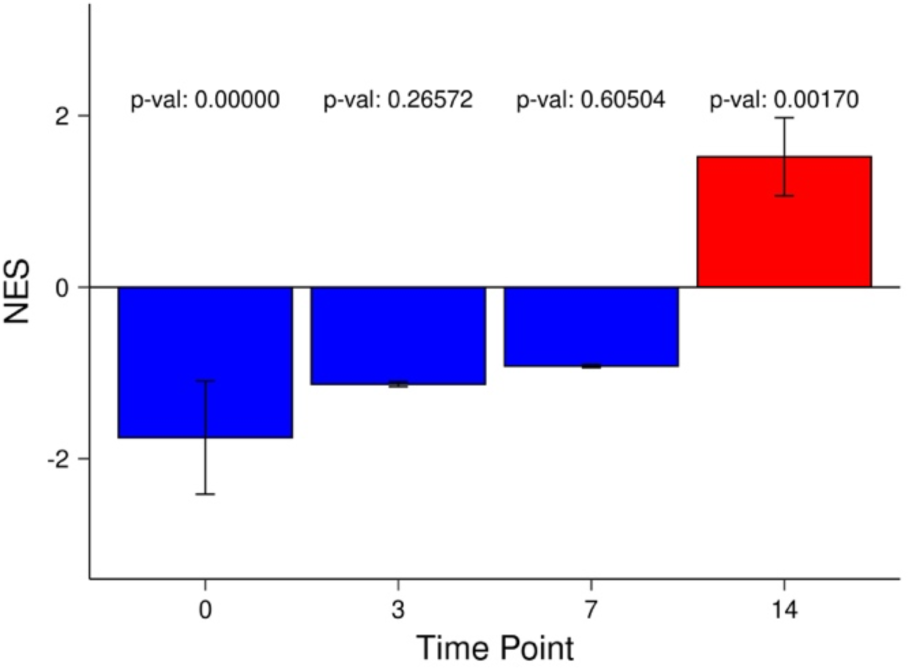
Enrichment of resistance signature along time points. Barplot showing the drug resistance signature gene set enrichment score at day 0, 3, 7, and 14 time points in the Oren et al. (2021) dataset. NES: Normalized Enrichment Score.

**Figure 1B:**
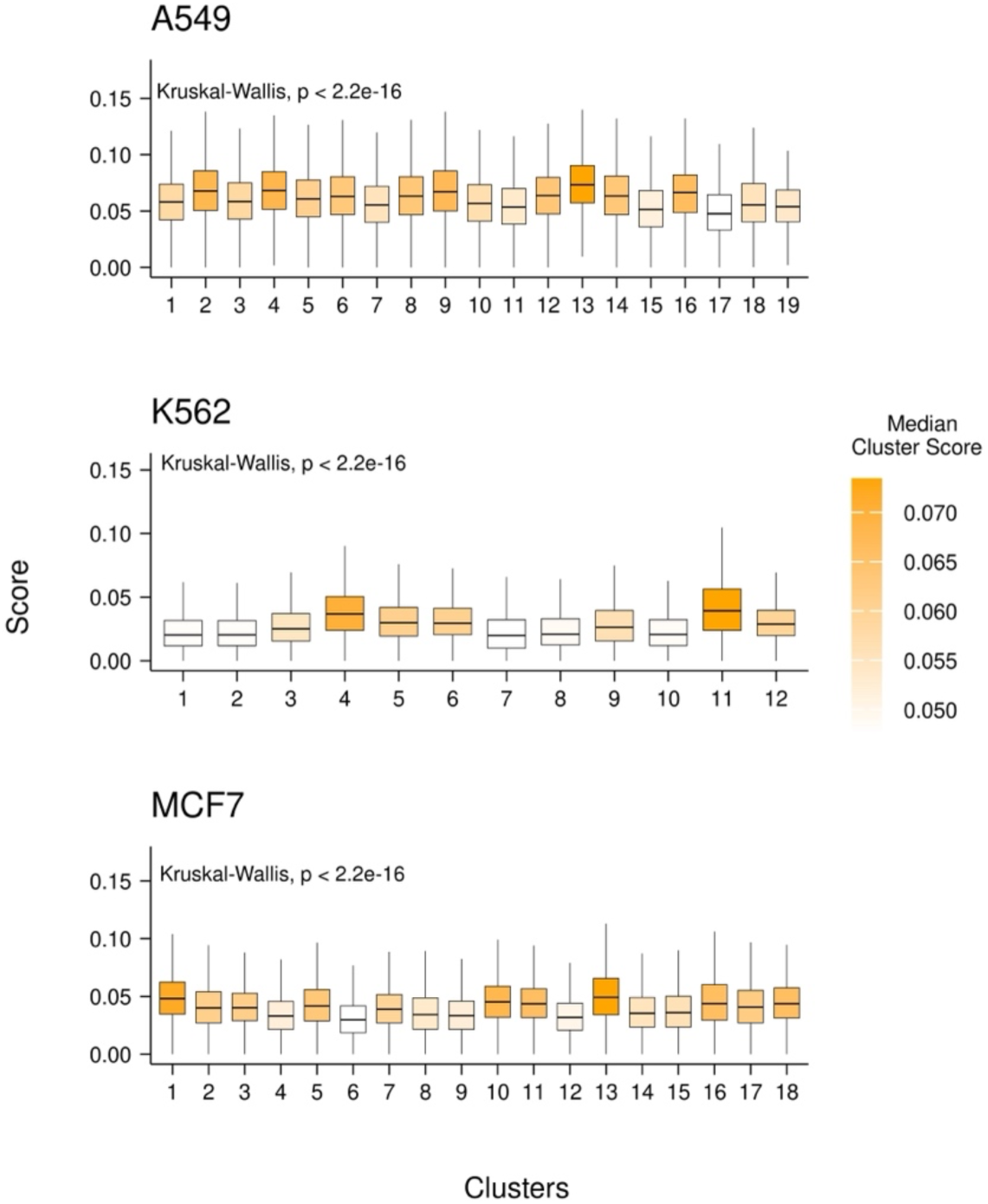
Resistance signature AUCell scores in cell line clusters. Raw AUCell score distributions for each cluster in each cell line. Each cell was scored with the drug resistance signature and the resulting distributions are shown as boxplots. The y-axis represents the raw AUCell score and each boxplot represents a transcriptional cluster. The color of a box plot represents the median raw AUCell score of that cluster.

**Figure 1C:**
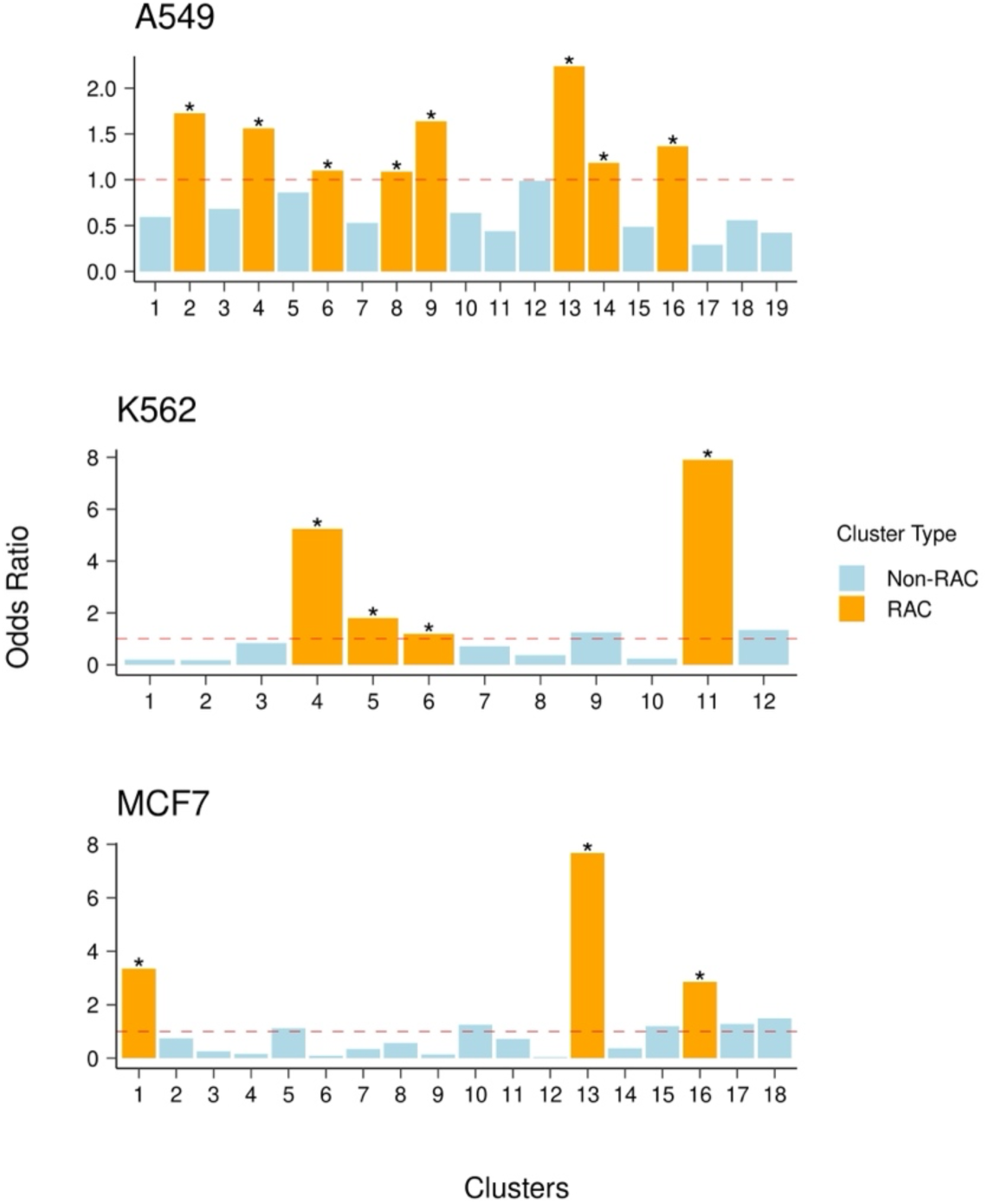
Resistance-active to inactive cell odds ratio of each cluster in each cell line. The y-axis represents the odds ratio and each bar represents a transcriptional cluster. Any cluster with an odds ratio greater than 1 and an adjusted p-value less than 0.05 is considered a Resistance Activated Cluster (RAC) and is colored orange, while the remaining clusters are non-RACs and are colored blue.

**Figure 1D:**
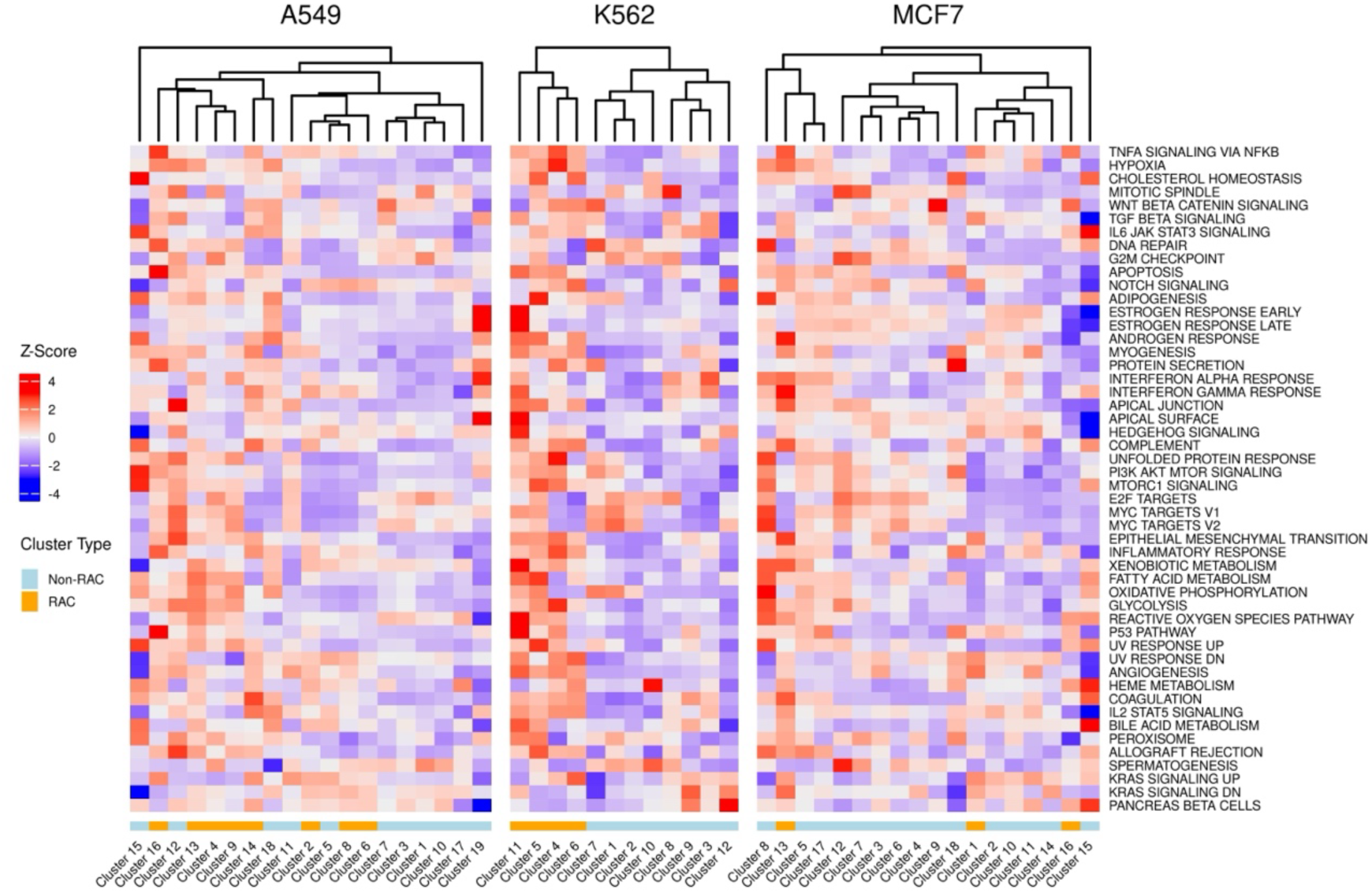
Z-score of each cancer hallmark’s average AUCell score of each cluster in each cell line. The rows represent the different hallmark genesets and the columns represent each transcriptional cluster, separately for each cell line. The colors in the bottom row signify which cluster is a Resistance Activated Cluster (RAC) (orange = RAC, blue = Non-RAC).

Next, we turned to a large scRNA-seq dataset containing drug-naive cells as well as cells 24 hours post-treatment, in three well-characterized cancer cell lines (A549, K562, MCF7) treated with 188 different compounds, belonging to 15 drug classes^11^. The variety of liquid and solid tumor types as well as multiple drug treatments in this dataset allows us to capture common aspects of drug response inherent to all cells regardless of cancer type. We downloaded the dataset of three single cell RNA-seq samples grouped into 49 transcriptional clusters by the original authors. Independently in each of the three cell lines, we assessed the relative expression of the resistance signature in each cluster using AUCell to get the raw signature score in each cell^16^ (see Methods). We observed that in each cell line, some of the transcriptional clusters had significantly higher resistance signature score distributions (Figure 1B). This suggests either pre-existence of the resistance signature expression or emergence within 24 hours of drug treatment.

Next, to identify the clusters or cell states that may represent the early stages of resistance, based on the AUCell score threshold (see Methods) to distinguish active from inactive cells for the resistance gene signature, we assessed the relative enrichment of resistance-active cells in each cluster relative to all other clusters of the same cell line (see Methods; Figure 1C). At an adjusted p-value threshold of 0.05 and odds ratio > 1, we identified multiple clusters in each cell line with markedly higher fractions of active cells compared to the baseline across the entire cell line. We termed them Resistance Activated Clusters (RAC).

Furthermore, we noticed that in each cell line some of the transcriptional clusters already had high levels of resistance signature expression even before the treatment, consistent with pre-existing resistance (Figure S1B). On the other hand, there are a number of emergent states (i.e. they only are considered active after the drug treatment suggesting acquisition of resistance) ranging from 2 to 5 states across the cell lines. These data clearly indicate that multiple cell states in pre-treatment and early drug response stages express resistance programs similar to those observed in long-term resistant samples.

### Resistance activated clusters are enriched for resistance pathways conserved across cell lines

Having identified RACs in each cell line, we aimed to functionally characterize them. We first scored all cells in each cell line with cancer hallmark signatures (from MsigDB^17^) using AUCell and plotted the mean score of each hallmark for each cluster (Methods; Figure 1D).

We observed that in two of the three cell lines, most of the RACs hierarchially clustering together, with some diversity in hallmark activity across RACs within each cell line, consistent with previously reported distinct, functionally diverged, resistant states^8^. Notably, in each cell line, there are RACs with both high enrichment and low enrichment of cell cycling signatures, consistent with previously noted cycling variability among resistance states^9^. We repeated this analysis with additional genesets, known as intratumor heterogeneity (ITH) meta-programs –– transcriptional programs that are commonly expressed across malignant cells in patient tumors^18^. The results of this analysis showed similar enrichment as the cancer hallmark analysis such as variable cycling and EMT expression (Figure S1C).

As a complement to the above analysis, in each cell line, we determined the differentially expressed genes between all RAC cells and all non-RAC cells using the Seurat FindAllMarkers function (Table S1; Methods), and assessed these genes for enrichment of Cancer Hallmarks, ITH meta-programs, and various GO biological processes and pathways (Methods; Figure S1D). Across all three cell lines, we observed enrichment of drug resistance-associated pathways such as EMT which previously has been shown to contribute to the development of drug resistance^19^. Other drug resistance-associated pathways enriched in the RAC signatures include TNFa and hypoxia, both associated with drug resistance in cancer^15,20^. The enrichment of these pathways further suggests that RAC cells may represent the early stages of drug resistant cell states.

Encouraged by the expression of resistance-associated programs among RACs within each cell line, we next aimed to identify pan-cell line resistance signatures. We hierarchically clustered the RACs across all three cell lines based on a gene expression similarity metric (see Methods) which yielded three groups (Figure S2A), each spanning correlated RACs from at least two of the three cell lines. We refer to the three pan-cell line RAC groups as superclusters (Figure 2A); superclusters 1 and 3 were represented by all three cell lines, while supercluster 2 was represented by two cell lines. To further probe the pan-cell line function of each supercluster, for each supercluster independently, we constructed two gene sets by combining the genes that are upregulated or downregulated in all three cell line-specific component clusters belonging to the supercluster (Methods; Table S1).

**Figure 2A:**
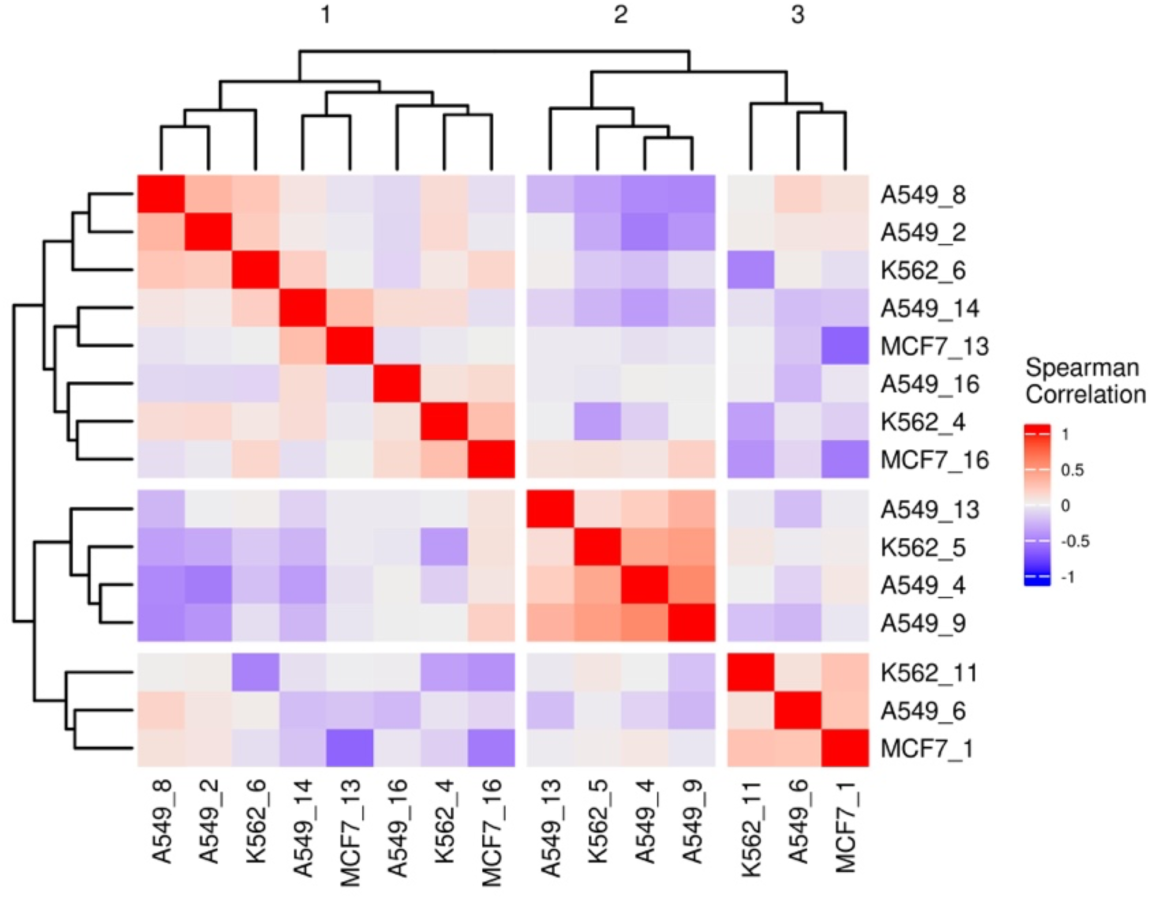
Correlations of RACs Differential Mean Expression of Most Variable Genes. Heatmap displaying the hierarchical clusters of an all by all correlation matrix based on the Spearman correlation of the cluster differential mean expression vectors (see Methods).

**Figure 2B:**
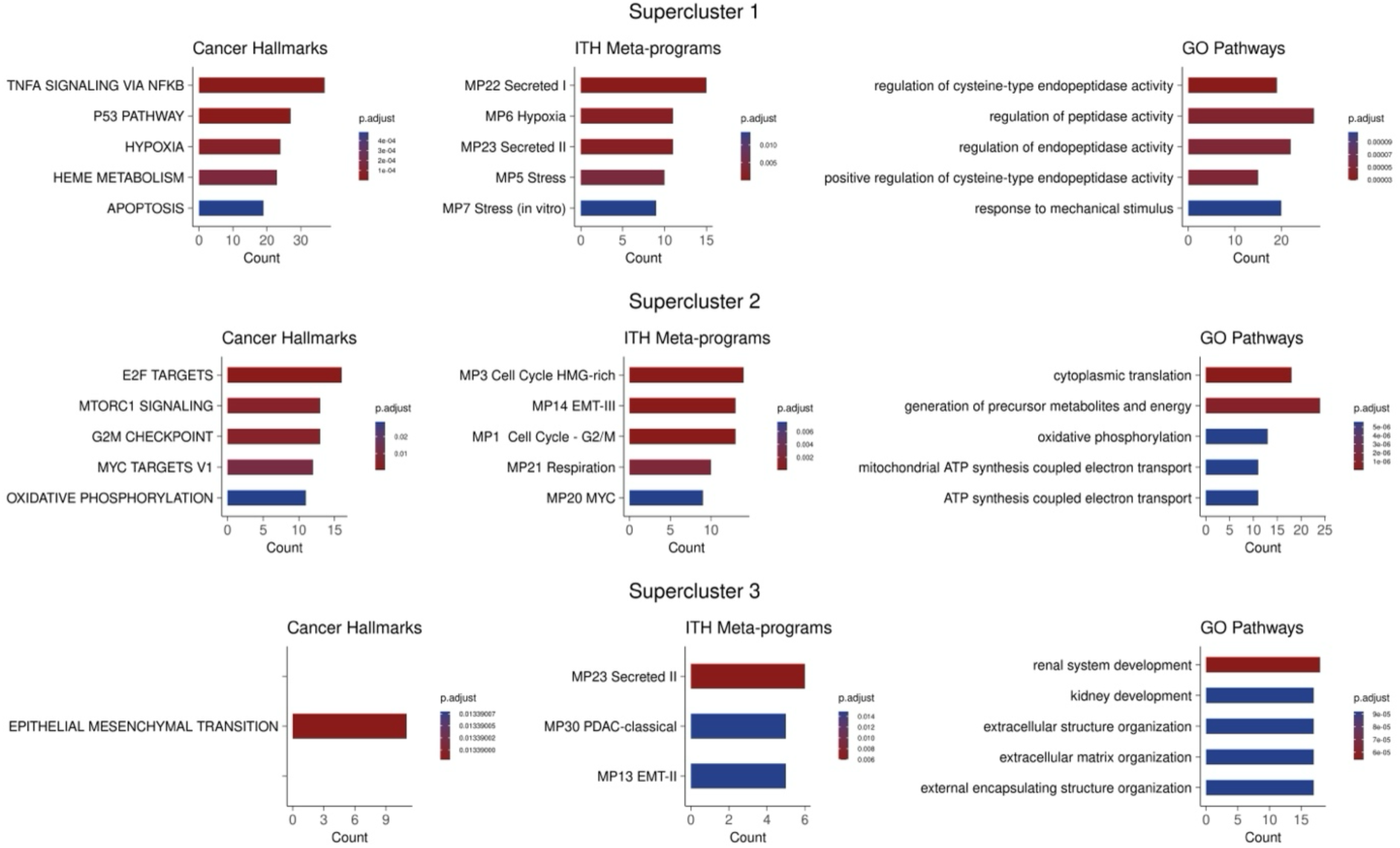
Supercluster Signatures Enrichment. The columns from left to right represent cancer hallmarks, ITH meta-programs, and GO pathways respectively. Within each column, the rows represent the three superclusters. Count values indicate the number of shared genes between the supercluster signatures and the enriched pathway.

**Figure 2C:**
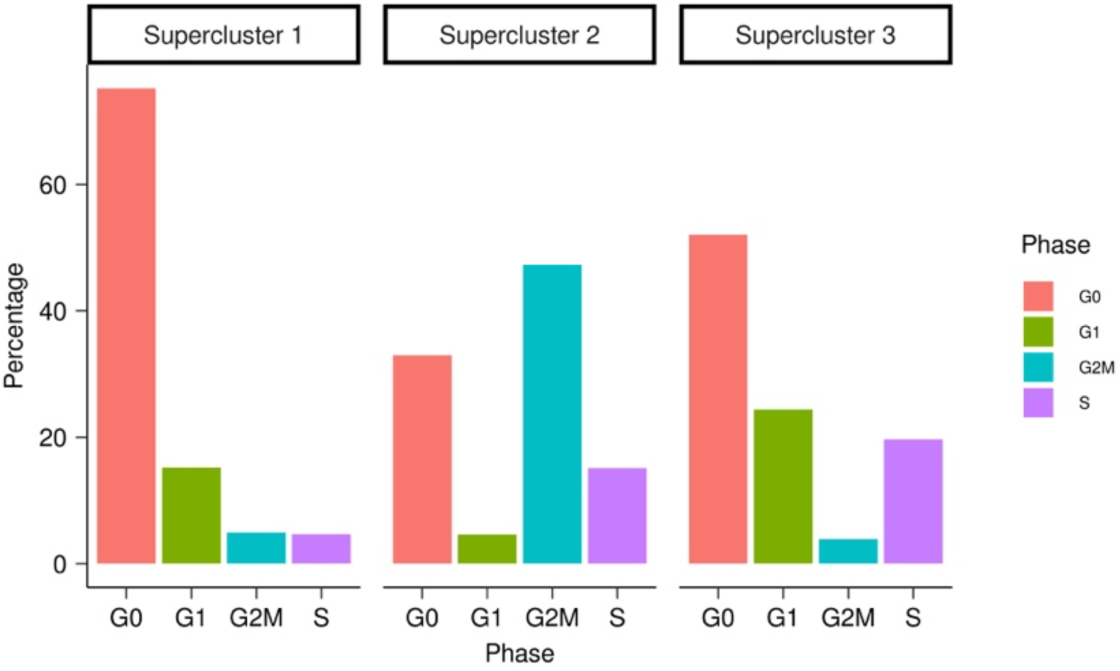
Superclusters Cell Cycle Phase Percentages. Each panel represents the percentage of total cells in each cell cycle phase for each supercluster. (G0 = red; G1 = green; G2M = blue; S = purple)

**Figure 2D:**
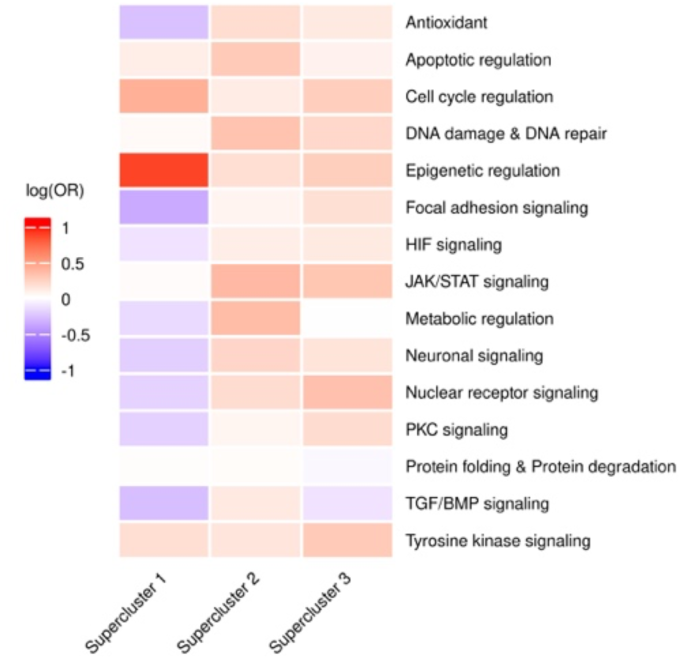
Drug Class Treated Cells Overrepresentation in Superclusters. Within each heatmap, the rows represent the various pathways targeted by the drug treatments and the columns represent each supercluster.

**Figure 2E:**
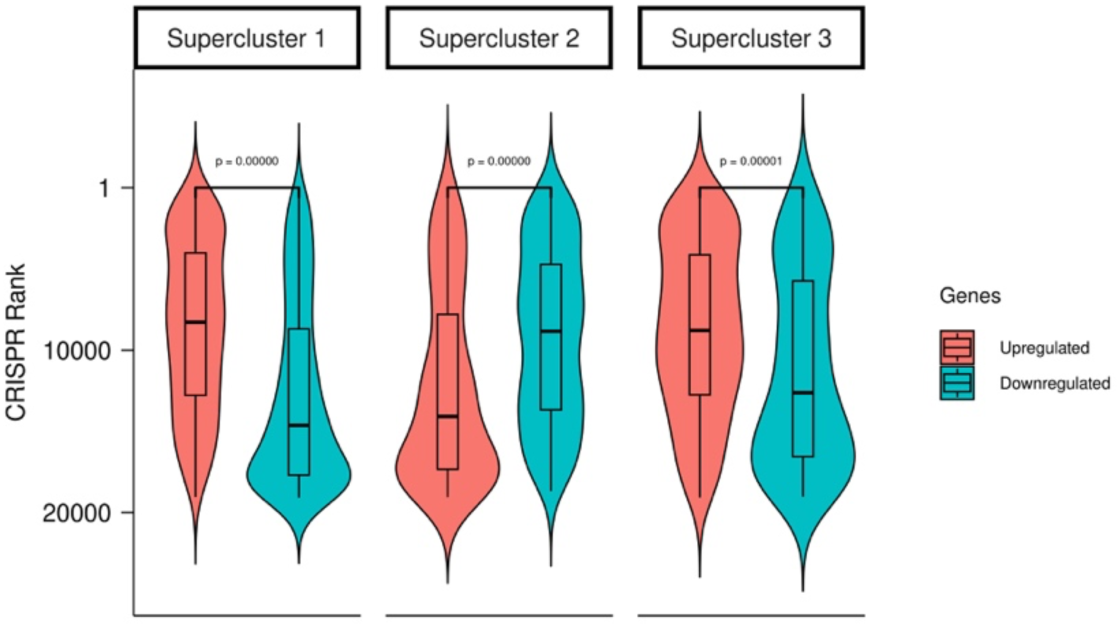
Distributions for CRISPR ranks for each supercluster up and downregulated gene (red = up; blue = down).

Functional enrichment analysis of the upregulated gene sets showed both distinct and shared enriched transcriptional programs across the three superclusters (Methods; Figure 2B). Notably, supercluster 1 is enriched for hypoxia and TNFa, which have been implicated in therapy resistance in multiple cancer types^20^. Hypoxia is known to directly cause drug resistance in cancer by decreasing drugs’ cytotoxic ability^15^, and hypoxia indicates increased cellular plasticity enabling these cells to survive drug treatment through adaptive responses^21^. Supercluster 2 was enriched for MYC activity which has been noted to contribute greatly to drug resistance^22,23^. The supercluster 2 signature also characterizes a change in metabolism associated with resistance^24^ through the enrichment of respiration and oxidative phosphorylation pathways. Finally, the enrichment observed in supercluster 3 included EMT and secretion pathways, both of which are assocated with resistance in various contexts^14,25^.

The pathways enriched in the supercluster 2 signature also included multiple cell cycling pathways indicating a cycling resistant state. This was further confirmed as a notable difference among the superclusters as the varying proportions of cell cycle phases (Figure 2C). This suggests variable proliferation rates across resistant states, previously attributed to differences in metabolism^9^. Specifically, supercluster 1, exhibiting the largest fraction of cells in G0 phase, may suggest a drug-resistant quiescent state^26^.

Repeating functional enrichment analysis of the downregulated gene sets in each supercluster resulted in very few pathways. Of the three superclusers, only supercluster 1 and 3 displayed any significant enrichment of functional pathways (Figure S2B). Among these pathways were mostly cell cycling pathways further indicating that supercluster 1 and 3 downregulate cycling pathways and exhibit a more queiscent state, a common feature of resistant states^27^.

Recall that our primary dataset includes responses to 188 drugs from 15 classes of drugs. We therefore assessed the associations of the superclusters with specific drug classes. To do this, for each supercluster, we calculated the enrichment of cells treated with each drug class in the RACs of each supercluster (Methods). We observed that supercluster 1 is largely composed of cells treated with drugs targeting epigenetic regulation, while supercluster 2 and 3 contain cells that are treated with a wider variety of drug classes (Figure 2D), suggesting a degree of specificity for a classes of drug to induce certain resistance state. These findings lead us to believe that some drugs may induce the same resistance state regardless of the cancer type, meaning that the transcriptional response is largely dictated by the treatment rather than the cancer type.

### CRISPR KO of genes driving resistance states sensitize the cells to the drug

Here, to assess broader validity of the resistance gene signatures, we experimentally tested if the genes defining the superclusters predictably impact resistance in a novel context. To do this we selected a cancer type and drug not present in any of the data analyzed to this point, and experimentally assessed whether knocking out the genes that characterize each supercluster increased sensitivity to cytotoxic drugs. To this end, we performed a genome-wide CRISPR knockout screen in an ovarian cancer derived cell line (OVCAR8) and quantified the effect of individual gene knockouts on the cell viability following treatment with a CHK1/2 checkpoint kinase inhibitor, Prexasertib, for 10 generations (see Methods). We ranked the genes based on the increase in drug sensitivity upon KO; higher ranking genes in this list are likely to mediate drug resistance. Encouragingly, we found that in supercluster 1 and 3, the upregulated genes have significantly higher CRISPR-based ranks than the downregulated genes. (p-value ∼ 0 for both; Figure 2E). The finding that the upregulated genes of supercluster 1 and 3 score better in the Prexasertib screen is consistent with the mechanisms of action of the drug, viz., replication catastrophe^28^. Due to low cell cycling of supercluster 1 and 3, cells occupying this state would therefore be naturally more resistant to this treatment, and accordingly, KO of genes associated with this state are expected to increase sensitivity to the drug. In contrast, cells with active supercluster 2 gene signature are expected to be more sensitive to Prexasertib treatment due to their higher expression of G2M checkpoint leading to susceptibility to replication catastrophe and cell death. And accordingly KO of genes associated with this state make the cell less sensitive to the drug, as reflected in the CRISPR screen (Figure 2E). This analysis suggests that disruption of resistance-associated transcriptional states in early response could contribute to drug tolerance.

### Conserved resistance states have clinical relevance

Having identified, and validated the RAC supercluster gene signatures, we next assessed the clinical relevance of each cell line’s global RAC signatures as well as the signatures of the three superclusters. First, for each cell line, we assessed the association of the cell line’s global RAC signature with patient survival in the matching cancer type in TCGA, i.e., A549 RAC signatures in lung cancer, K562 RAC signatures in blood cancer, and MCF7 RAC signatures in breast cancer. We scored the samples with each global RAC signature using ssGSEA^29^ and performed a Cox regression analysis on overall survival while controlling for age, sex, and tumor purity (Methods). In all three cancer types, high expression of the global RAC signature is associated with worse survival with hazard ratios of 2.1, 2.5, and 2.0, respectively (Figure 3A).

**Figure 3A:**
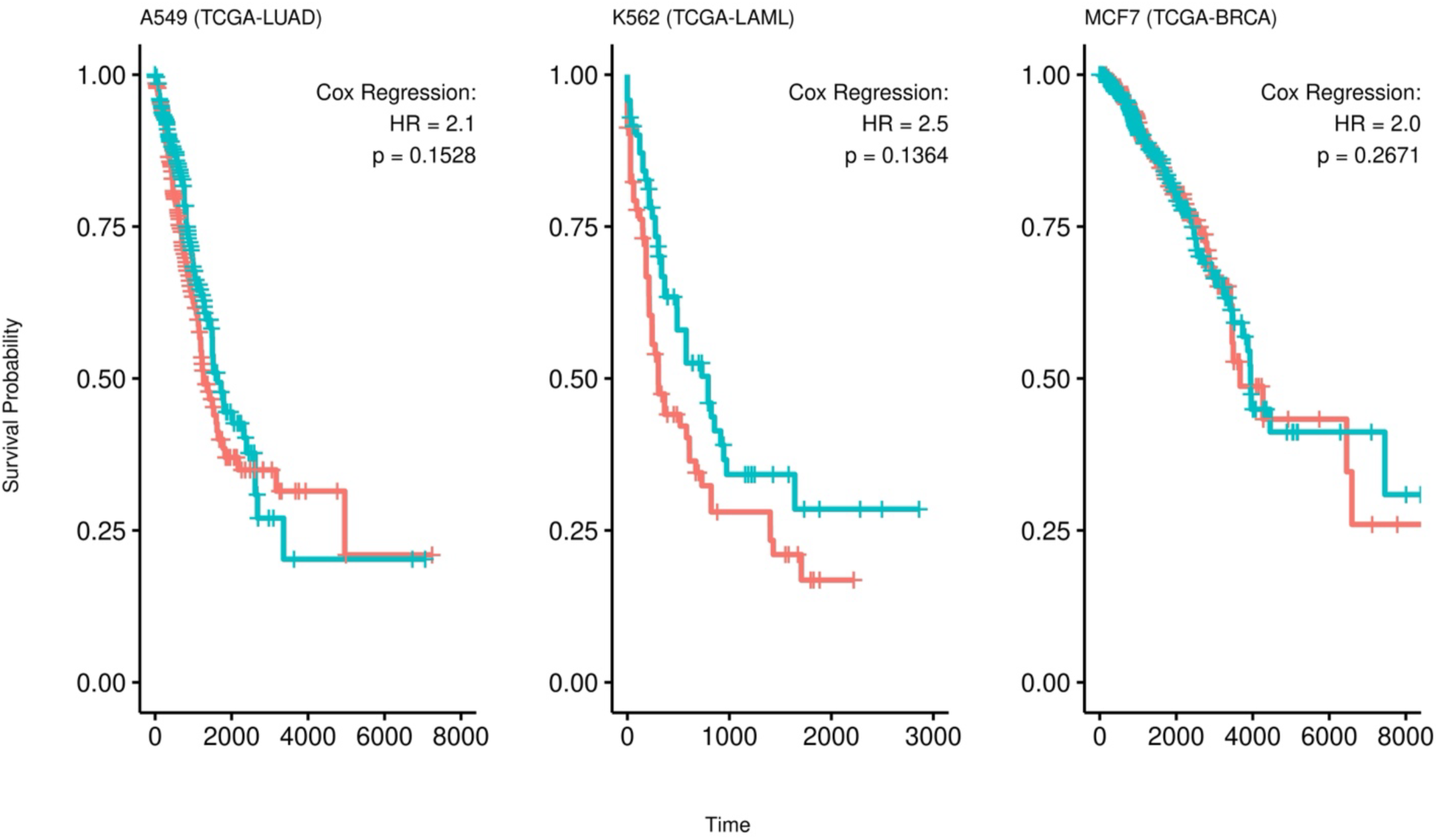
Global RAC Signatures are associated with overall survival in the corresponding cancer. Kaplan-Meier plots for the survival analysis of each RAC Type signature from each cell line. The patients are stratified into 2 groups representing their level of expression for the given cell line signature (red = high, blue = low). The HR and p-values are obtained from Cox regression.

**Figure 3B:**
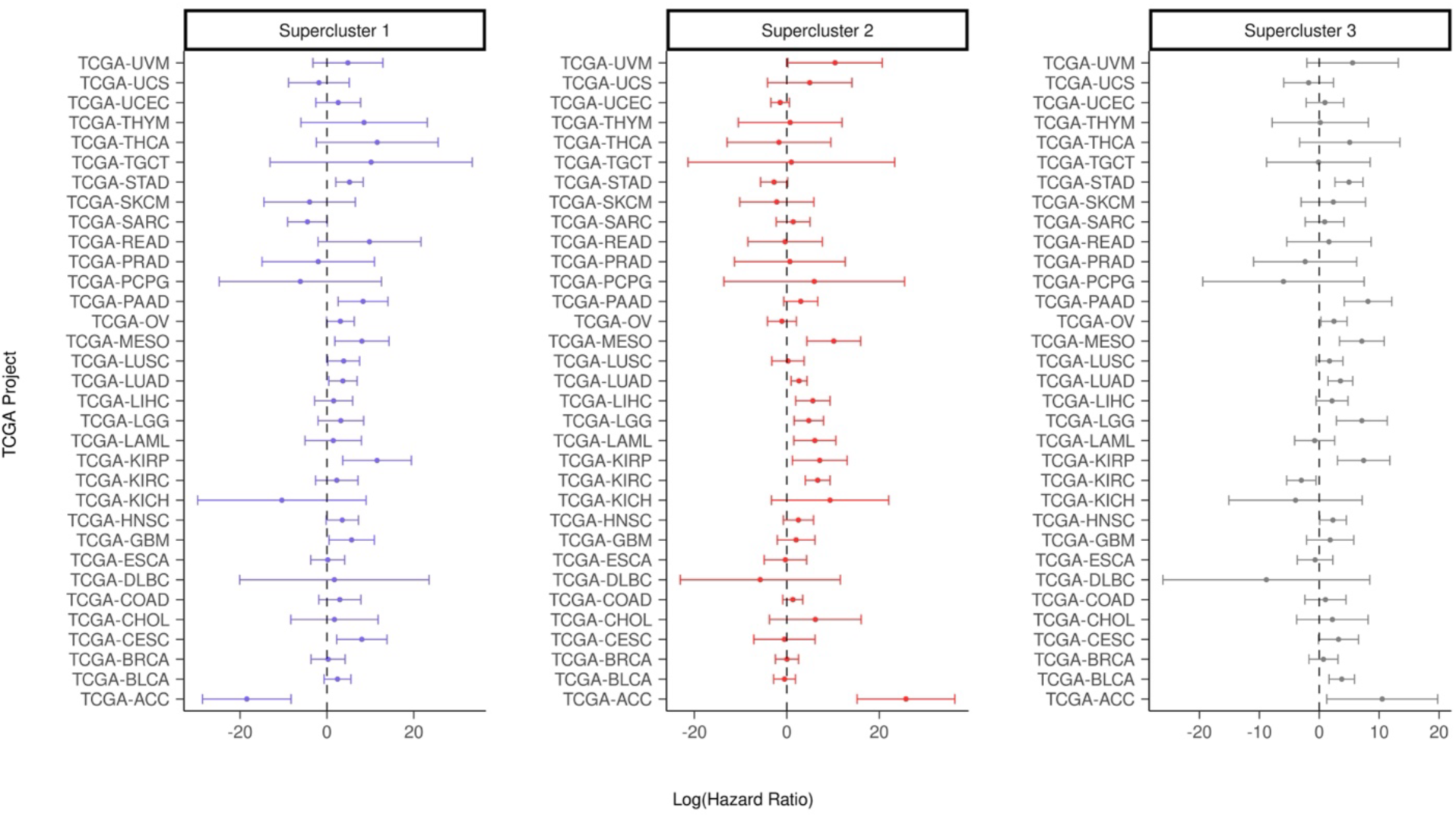
Hazard Ratios for Supercluster Signatures Across TCGA Cancer Types with 95% confidence interval of log hazard ratios for all supercluster signatures in all 33 TCGA projects.

**Figure 3C:**
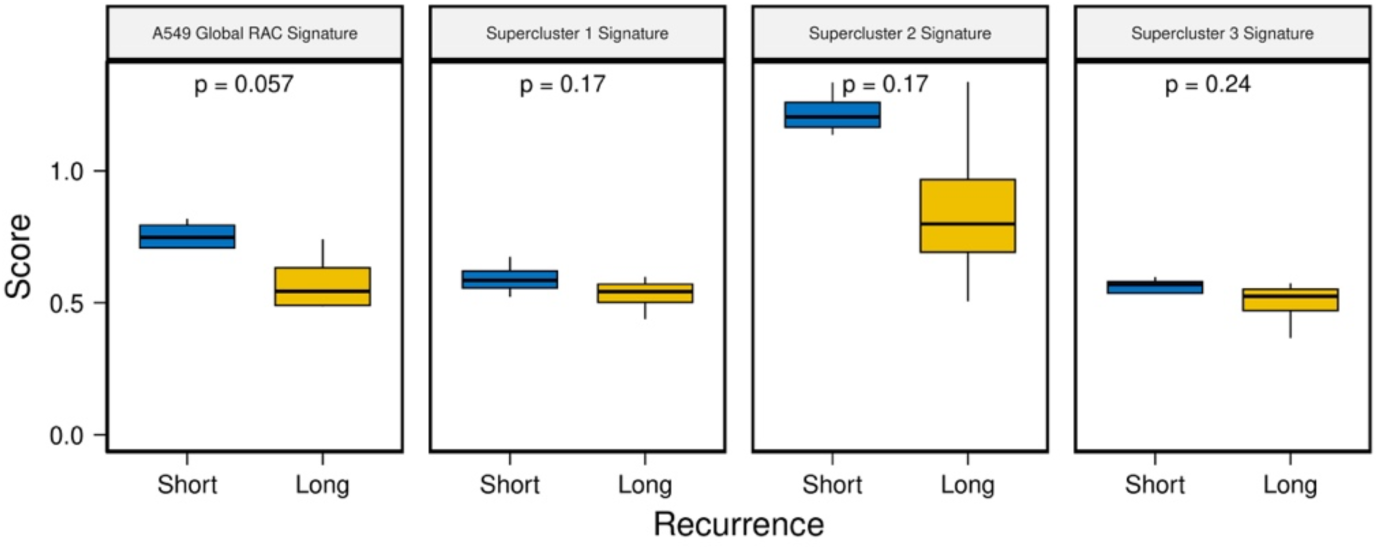
Pre-treatment RAC signatures scores in lung cancer patients before tyrosine kinase inhibitor treatment. The y-axis represents the ssGSEA score and the x-axis stratifies patients by recurrence time (blue = short term recurrence, yellow = long term recurrence).

**Figure 3D:**
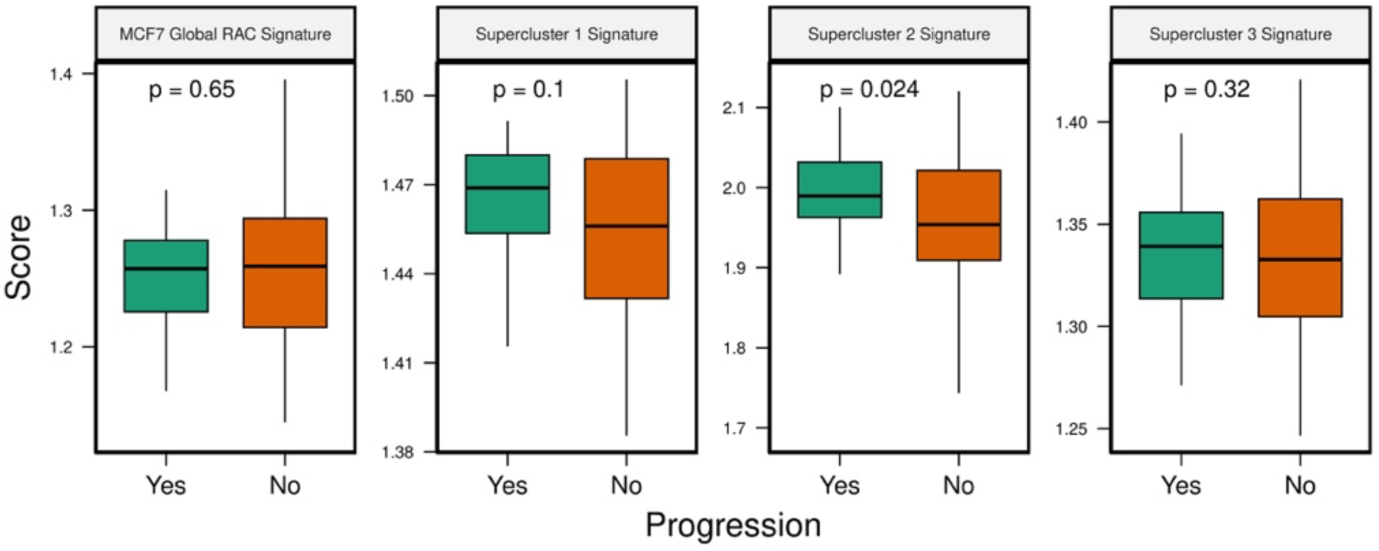
RAC signature scores in premalignant lesions. The y-axis represents the ssGSEA score and the x-axis stratifies patients by progression status (green = progression, orange = no progression)

**Figure 3E:**
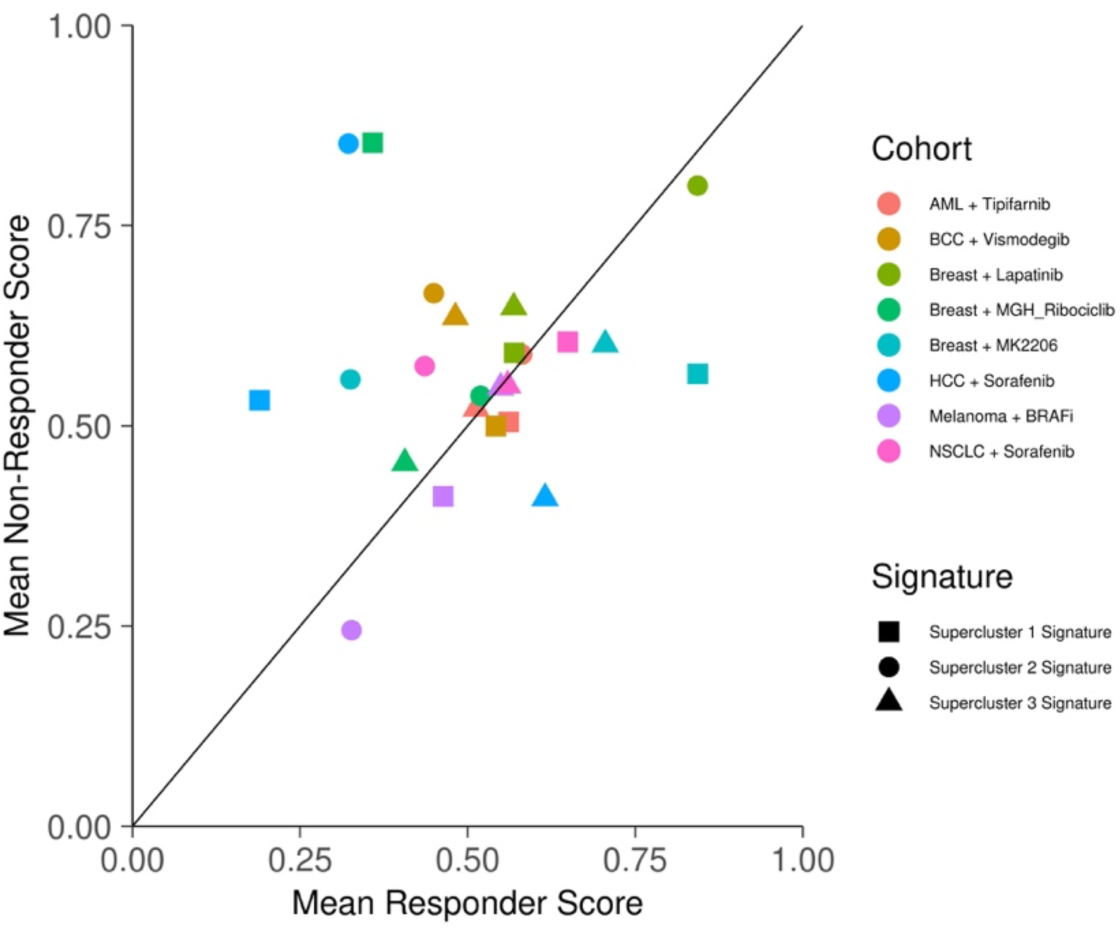
Mean supercluster signature scores for responders and non-responders across cancer treatment cohorts (cohort = drug x cancer combo).

These results indicate that the signatures associated with resistance derived from early drug response are prognostic of overall survival in all of the three cancer types. Since superclusters are representative of multiple cancer types, we repeated the above analysis with the supercluster signatures across all TCGA cohorts. Although we observed, not surprisingly, substantial variability across cancer types, the distribution of log-transformed hazard ratios are significantly greater than 0 with p-values of 0.0176, 0.0016, and 0.0082, respectively. (One sample t-test) (Figure 3B).

Given the RAC and supercluster signatures’ association with patient survival, we next assessed their ability to predict therapy response. Toward this, first, we acquired a dataset that contains pre-treatment tumor RNA-seq samples from 8 patients with EGFR-mutant lung cancer treated with the tyrosine kinase inhibitor, Osimertinib^30^. All patients had relapsed after initial response and were classified into those with short relapse time (4 patients, < 8.6 months) and those with long relapse time (4 patients; >= 12 months). We scored each patient’s pre-treatment sample with the A549 global RAC signature (cancer type matched) as well as with the three supercluster signatures using ssGSEA. We observed that the patients with short relapse times had higher resistance scores for the A549 global RAC signature and all supercluster signatures; p-values are likely not significant due to the small sample size (n = 8, Figure 3C). This suggests pre-existing resistance mechanisms in short-term responder patients.

We next assessed the extent to which supercluster signature may be predictive of therapy response across cancer and drug types. The data we used consisted of bulk RNA sequencing and microarray data spanning 6 cancer types and 7 drug treatments, where in each study the patients were classified into responders and non-responders^31^ (Table S3). We found that globally across all cohorts and supercluster signatures, on average the non-responders scored higher for the supercluster signatures, although the significance was marginal (one-sided t-test p-value = 0.08; Figure 3E).

Lastly, we assessed whether the supercluster signatures are associated with progression of pre-malignant lesions. Toward this, we obtained a dataset (RAHBT97 cohort) from the HTAN database^32,33^ consisting of bulk RNA sequencing of pre-malignant breast lesions that either progressed (31 samples) to cancer or not (66 samples). We then scored each pre-malignant sample with the MCF7 global RAC signature (cancer type matched) as well as with the three supercluster signatures, using ssGSEA. Comparing the of scores between samples that progressed to samples that did not, we observed significantly higher scores in the progression samples for the supercluster 2 signature and while in the other supercluster signatures the difference was not statistically significant, we observed the same trend (Figure 3D). Interestingly, we also observed a significant enrichment of genes differentially expression in progression samples in the signature for supercluster 2 (Figure S2C). These findings indicate that the resistance-associated signatures we generated can predict malignant progression.

Overall, our results suggest that our cancer-specific RAC as well as pan-cancer supercluster resistance signatures are prognostic of patient survival, tumor progression, and therapy response across cancers and therapy types.

### Supercluster resistance mechanisms are evolutionarily conserved

Having identified common mechanisms of resistance across the three cancer cell lines, and observing their prognostic utility across cancer types, we next assessed the extent to which these resistance-associated transcriptional responses represent inherent cellular response to chemical-induced stressors and therefore may be evolutionarily conserved across different species. To test this hypothesis, we first obtained a gene signature comprising the genes that are upregulated in the yeast *Candida auris* resistant to antifungal drugs^13^ and mapped them to their human orthologs using information from OMA Browser^34^ (Table S4; Methods). Next, we scored all cells in the early-response dataset using AUCell and compared the score distributions across the various supercluster cells and the non-RAC cells (Methods). In supercluster 2 and 3, we saw significantly higher signature scores for the ortholog yeast resistance geneset compared to all non-RAC cells with effect sizes of 0.58 and 0.70, respectively (Cohen’s d: measures difference between the standardized means of two distributions). This suggests that supercluster 2 and 3 represent conserved mechanisms of drug response between yeast and human (Figure 4A).

**Figure 4A:**
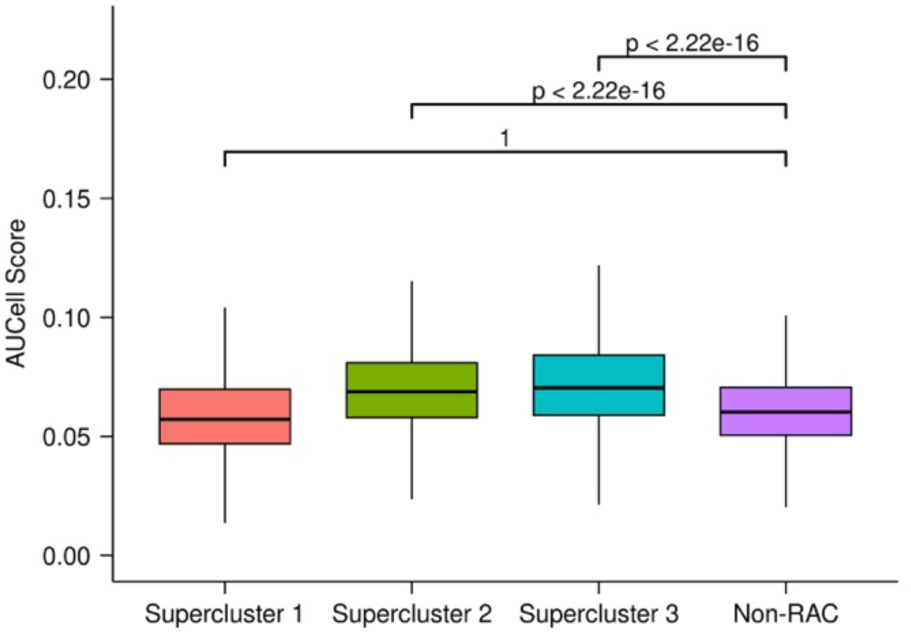
Yeast antifungal resistance gene signature scores in human resistance and non-resistance clusters. Each distribution contains the scores for every cell in each supercluster or non-RAC clusters across the three cell lines.

**Figure 4B:**
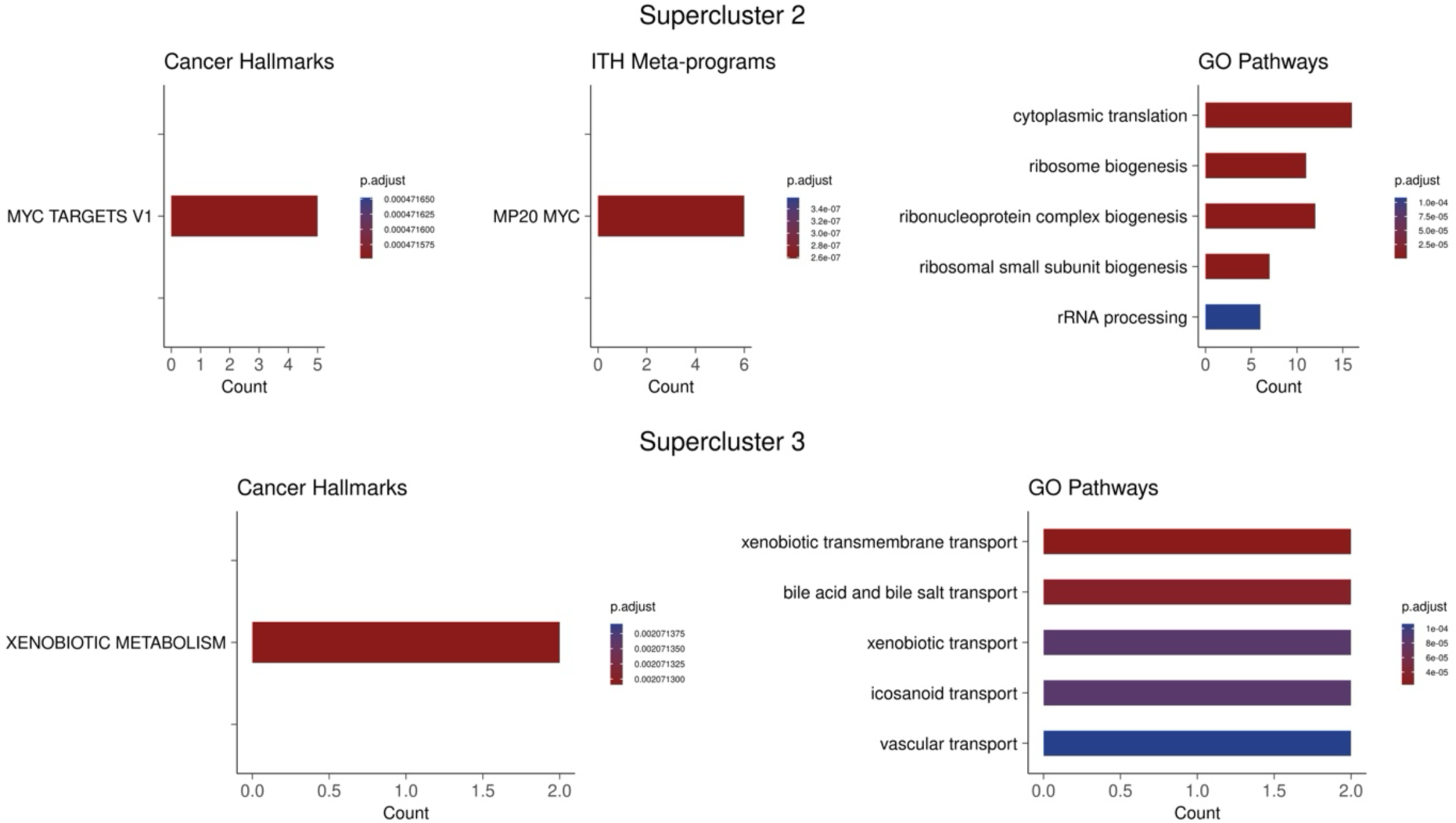
Pathway enrichment of shared genes between yeast resistance orthologs and supercluster 2 and 3. Count values indicate the number of shared genes between the genes of interest and the enriched pathway.

**Figure 4C:**
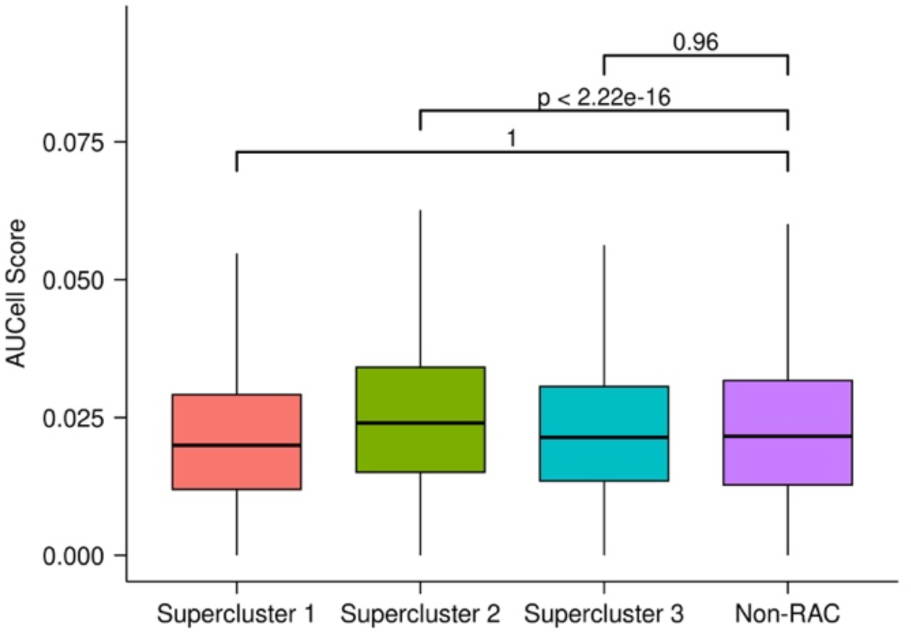
E. coli antimicrobial resistance gene signature scores in human resistance and non-resistance clusters. Each distribution contains the scores for every cell in each supercluster or non-RAC clusters across the three cell lines.

**Figure 4D:**
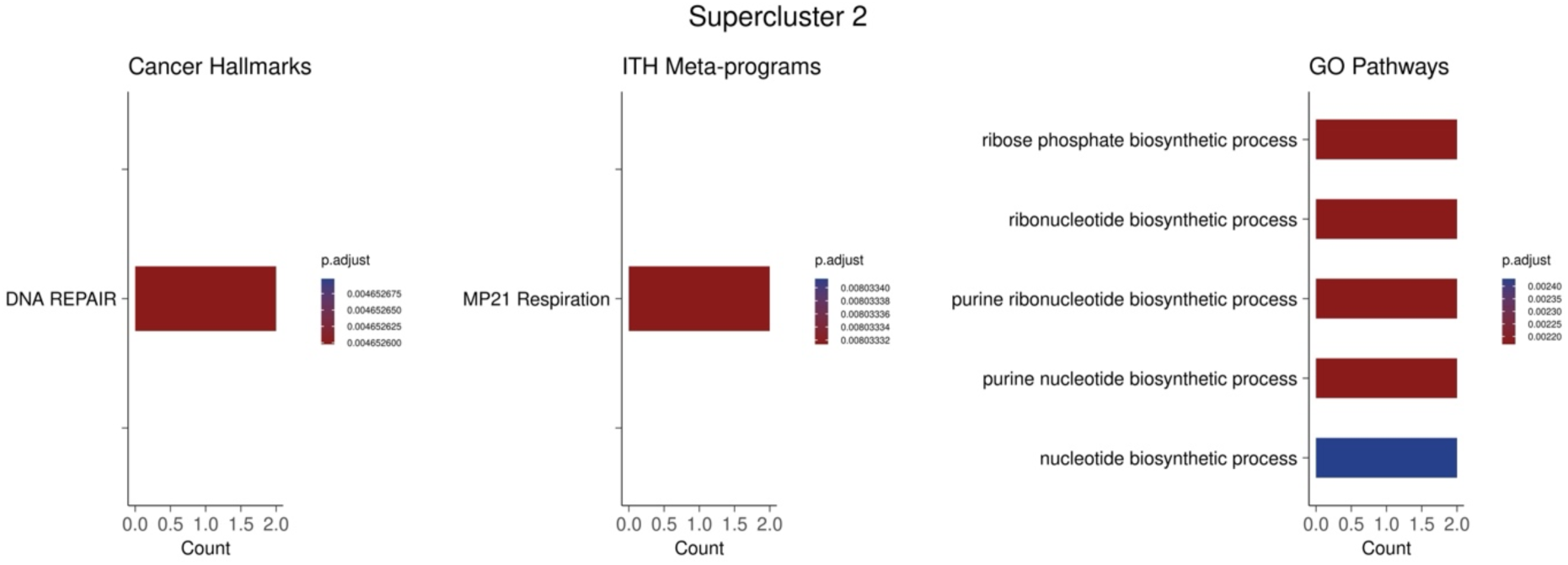
Pathway enrichment of shared genes between E. coli resistance orthologs and supercluster 3. Count values indicate the number of shared genes between the genes of interest and the enriched pathway.

To further probe into the underlying shared mechanisms, we performed functional enrichment on the genes that are common in both the antifungal resistant ortholog geneset and genes upregulated in supercluster 2 and supercluster 3 (Figure 4B). This analysis showed enrichment for MYC activity and ribosome biogenesis in the shared genes between supercluster 2 and the yeast orthologs, suggesting that these processes are conserved properties of cellular response to chemicals and may contribute to resistance. Increased MYC activity and ribosome biogenesis is consistent with previous literature as MYC is a master regulator of ribosome biogenesis^35,36^ and these processes have been shown to play a role in drug resistance in both yeast and in human cancer^22,37,38^. The pathways enriched between supercluster 3 and the yeast orthologs include xenobiotic metabolism and various transport pathways. Notably, the genes contirbuting to the enrichment of these pathways include ABCC2 and ABCC3 (*C. auris* ortholog: B9J08_002146). These genes code for ATP-binding cassette (ABC) transporters which have been shown to directly lead to multidrug resistance in both human cancer and yeast^39,40^. These finding suggests the conserved resistance mechanism between human cancer and yeast rely heavily on MYC and its downstream processes as well as drug efflux pumps.

Next, we compiled experimental data of drug-treated *E. coli* strains and obtained differentially upregulated genes in resistant relative to drug-naive samples for 9 antimicrobial drugs. With the 9 drug resistant upregulated genesets, we created a consensus antimicrobial resistance gene set (Table S4; Methods) and mapped these genes to their human orthologs and as in the case of yeast, we scored all cells for the gene signature using AUCell. In supercluster 2, we saw significantly higher resistance scores compared to all non-RAC cells with an effect size of 0.16 (Figure 4C).

We next performed functional enrichment on the genes that are shared in both the antimicrobial resistant ortholog geneset and genes upregulated in supercluster 3 (Figure 4D). The pathways showing enrichment indicate the mechanisms that are evolutionarily conserved between *E. coli* and human cancer seem to stem mainly from increased DNA repair. This finding is consistent with previous literature as enhanced DNA repair mechanisms have been noted to contribute to resistance in both *E. coli* and human cancers^41,42^. Overall, these analyses suggest that supercluster 2 and supercluser 3 represent evolutionarily conserved mechanisms underlying drug resistance between human and yeast involving the known proto-onco gene MYC, its downstream processes like ribosome biogenesis, and drug efflux as well as supercluster 3 representing evolutionarily conserved mechanisms underlying drug resistance between human and *E. coli* involving enhanced DNA repair mechanisms.

## DISCUSSION

In this work, we have comprehensively assessed the extent to which the transcriptional programs that characterize drug-resistance in cancer cells either exist before drug treatment or are manifested early in the course of drug treatment and thus may represent inherent cellular response to cytotoxic stressors, some of which form the basis for long-term resistance. We find that many of the transcriptional programs that are observed in established resistance such as epithelial to mesenchymal transition^43^, MYC signaling^22,23^, and oxidative phosphorylation^24^ are indeed present just 24 hours after drug treatment. Furthermore, not only are these early manifestations of resistance programs evident in multiple cancer cell lines, but aspects of them exhibit deep evolutionary conservation in bacteria and yeast, suggesting a conserved inherent cellular response to cytotoxic stressors, ultimately linked to long-term survival of the cell. Previous studies have shown that genes with higher expression variability manifest as differentially expressed in disparate experiments and are enriched for stress response genes^44,45^. Ultimately, directed experiments are the only way to ascertain causality. Using a whole genome CRISPR knockout screen, we validated that the genes defining our identified early drug resistance states, when knocked out, render the cell more sensitive to the drug, in a manner consistent with the drug’s mechanism of action. Coupled with the prognostic nature of these signatures in pre-malignancy and overall survival, these signatures that were inferred from early response data may hint at the existence of cell states resistant to multiple stresses that cells have encountered across evolutionary time scales.

Consistent with previous reports^8^, our analysis reveals distinct modes of resistance. The resistant transcriptional states conserved across cancer types exhibit varying cell cycle states and association with distinct drug classes (Figure 2). These aspects of resistance are frequently outlined in the literature as responses to drug treatment^9,26^. Interestingly, in our analyses, these resistance-linked early response signatures even in pre-treatment tumor transcriptomes are associated with duration until relapse, overall survival, therapy response, and tumor progression, across multiple cancer and treatment types (Figure 3). Since these predictions are based on tumor data obtained at the time of diagnosis, and not after cancer chemotherapy, expression of the identified resistant states may be present before treatment and subsequently selected for upon treatment. On the other hand, multiple transcriptional states appear strictly post-treatment and may represent adaptive responses (Figure 1C, Figure S1B). Our findings corroborate recent reports on drug-tolerant persister (DTP) cells, which are rare, slow-cycling persister cells that transiently occur post-treatment and can revert to cycling upon release from treatment and become sensitive to the initial therapy^6,46^. However, it is still unclear whether DTPs are pre-existing states or the result of therapeutically induced transcriptional reprogramming^7^ which could be best investigated via lineage tracing. An additional confounder to consider is that rare populations of pre-existing resistant cells may go undetected in single cell profiling and may appear as emergent states post-treatment.

Although we observed expression of our resistance state signatures across multiple cancer types, the robustness of this signature could be improved. Our study derives the initial resistance signature by compiling just 6 datasets and the early response signature from 3 cell lines. By integrating additional datasets these signatures would become more generalizable and perhaps uncover additional resistance mechanisms not captured by the approach in this work.

The drug response data used in our study includes responses to a large array of drugs and drug classes and therefore the identified resistance-associated signatures are relevant across multiple drug classes^11^. One of the prominent mechanisms underlying multidrug resistance (MDR) involves ATP-binding cassette (ABC) transporters. We found that all three cell type-specific global RAC, supercluster 1, and supercluster 3 gene signatures include ABC transporter genes (ABCA1, ABCA12, ABCB1, ABCC2, ABCC3, ABCC13, and ABCG1) suggesting their importance in resistance across cell lines. While it has been shown that the expression of one ABC transporter, P-glycoprotein 1 (Pgp, ABCB1), can be induced by treatment^47^, it is unclear to what extent Pgp plays a role in the resistance characterized in this work.

Novel treatment strategies to modulate resistance programs in cancer are currently being investigated^2^. Among them are targeting biomarkers associated with drug resistant cancer cells, and the supercluster signatures identified in this work may represent potential avenues of further investigation^48^, as supported by our CRISPR screen. One future angle of investigation may involve assessing whether the genes underlying the enriched pathways are the same or different across different cancer types. This would allow for cancer type-specific treatment approaches for common resistance pathways. Although much more research is needed to fully understand the mechanisms of resistance and how to inhibit them, this work contributes to the growing knowledge of the molecular characteristics of resistant transcriptional states shortly after treatment which may inform novel adjuvant treatment strategies to overcome resistance.

## RESOURCE AVAILABILITY

### Lead contact

Further information and requests for resources should be directed to and will be fulfilled by the lead contact, Sridhar Hannenhalli.

### Materials availability

This study did not generate new materials.

### Data and code availability

- All data needed to evaluate the conclusions in the paper are present in the paper and/or the Supplementary Materials. Public datasets used in this study are available in referenced papers. Source code and analysis scripts are available on GitHub (https://github.com/coleruoff/drug_treatment)

## ACKNOWLEDGMENTS

Authors thank members of Hannenhalli lab and Gottesman lab for helpful discussion.

## AUTHOR CONTRIBUTIONS

Conceptualization, CR, SH, VG methodology, CR, AM, PM, VG Investigation, CR, AM, PM; writing—original draft, C.R. writing—review & editing, CR, SH, AM, PM, VG, MG; funding acquisition, SH, MG; resources, SH, MG; supervision, SH, MG

## DECLARATION OF GENERATIVE AI AND AI-ASSISTED TECHNOLOGIES

The authors report no use of generative AI or AI-assisted technologies.

## SUPPLEMENTAL INFORMATION

**Document S1. Figures S1-S2**

**Table S1. Resistance Signatures**

**Table S2. CRISPR Gene Ranks**

**Table S3. Responder/Non-responder Scores**

**Table S4. Ortholog Gene Sets**

## STAR★METHODS

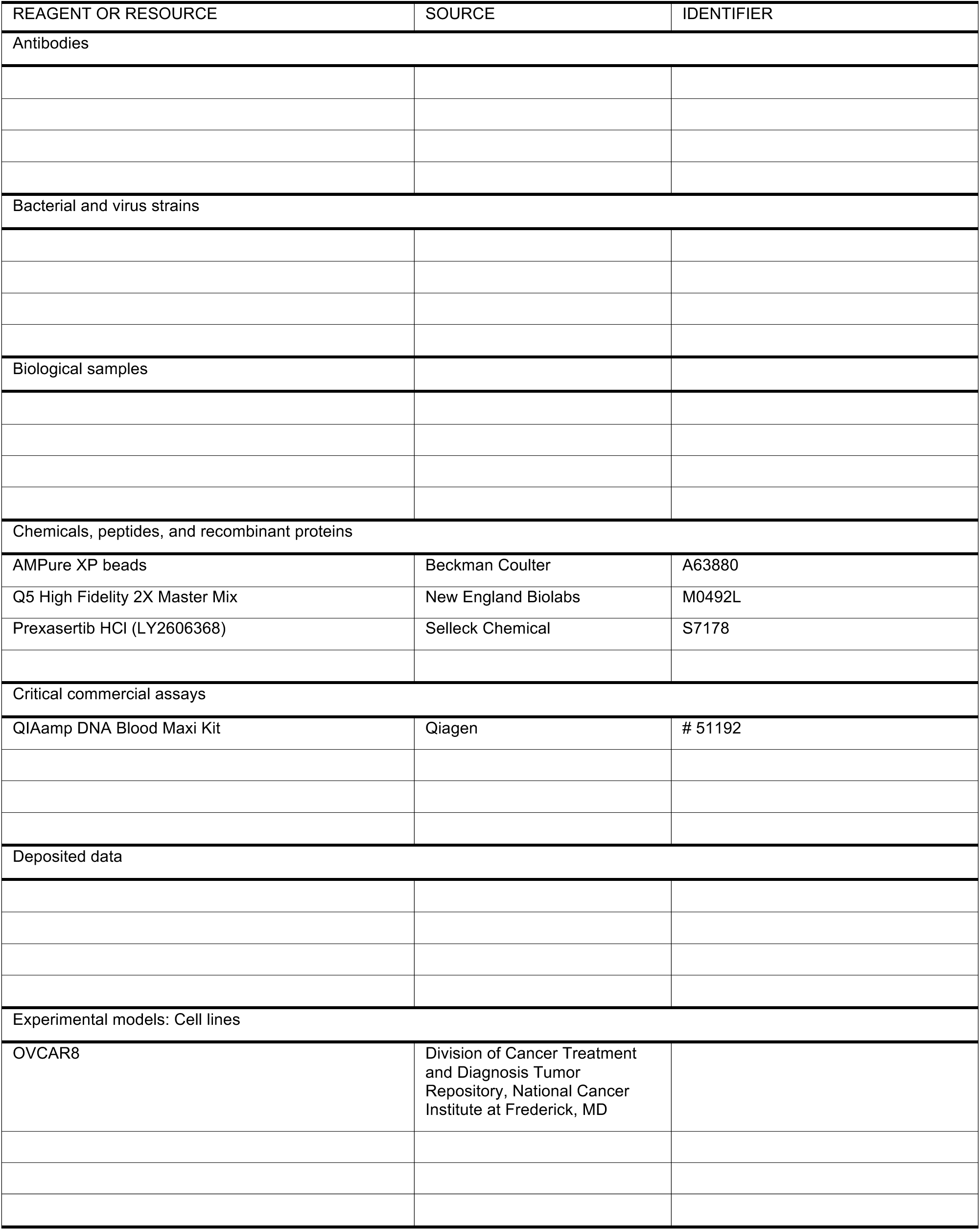
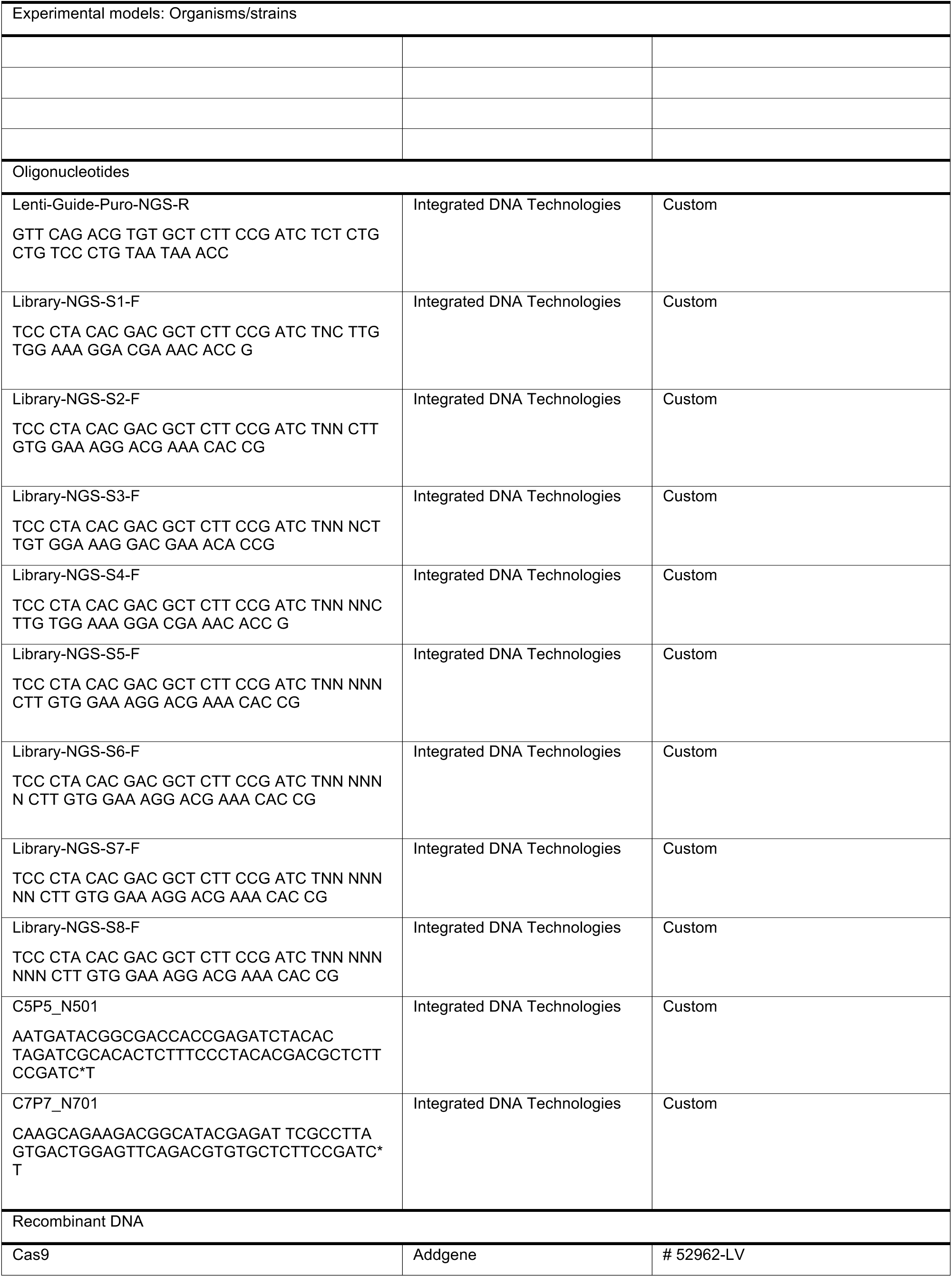
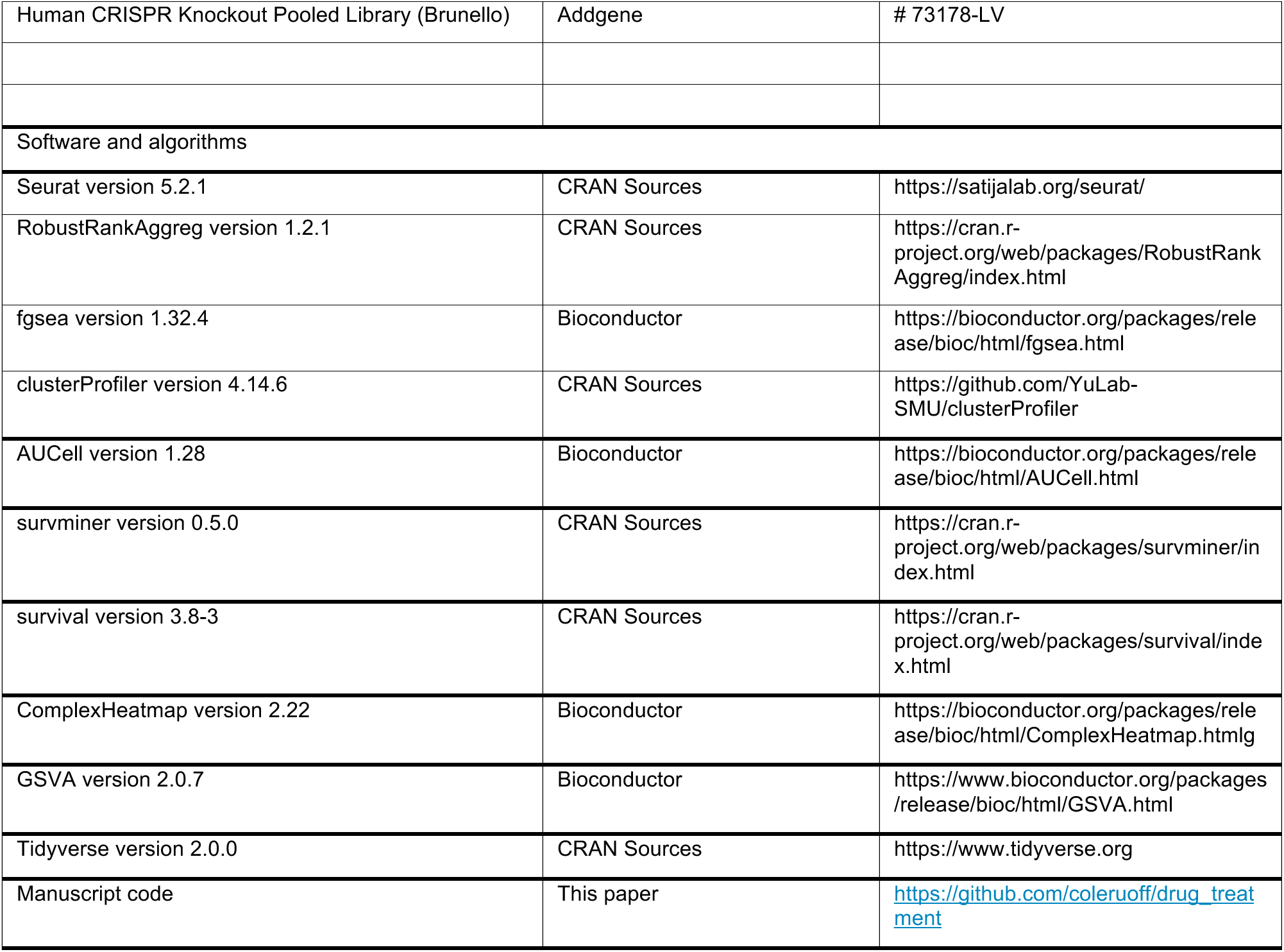
KEY RESOURCES TABLE.

## METHODS

### Single Cell Datasets

For the Oren et al. dataset, we first downloaded the raw count matrices from the publicly available repository referenced in the original publication. Within each cell line, we filtered out the cells with less than 200 and greater than 2500 features then normalized the data using log transformation with a scale factor of 10,000.

For the Srivatsan et al. dataset, we downloaded the filtered count matrices from the publicly available repository referenced in the original publication. Within each cell line, we further filtered out the genes that are expressed in less than 100 cells to retain genes driving biological processes. This was followed by normalization of the data using log transformation with a scale factor of 10,000. Pre– and post-treated cells were processed identically, removing any treatment stage batch effects.

### Creating resistance signature

We created a drug resistance gene signature (72 genes) that consists of the aggregation of ranked lists from 6 drug treatment experiments in multiple cell lines (COLO829, A375, HT29, MMACSF, EFM192A, and BT474)^9^. In these experiments, the cell lines were exposed to the drugs for 10 days and at day 10 the surviving cells were sequenced. We repeated the single cell processing approach as detailed in the publication’s methods section then for each cell line, we performed differential expression between day 0 and day 10 using the FindAllMarkers() function. We next selected all genes that were upregulated at day 10 in each cell line (average log2FC > 0 and adjusted p-value < 0.05) and sorted all six gene lists by log2FC. Finally, we formed a single ranked list using Robust Rank Aggregation (RRA)^49^ and further filtered to retain all genes with a adjusted rank p-value < 0.05.

We validated the gene signature in time-course data of drug resistance development (Oren at al. REF)-PC9 lung cancer cells were treated with Osimertinib and single cell transcriptomic profiling was done at days 0, 3, 7, and 14. For each timepoint in the Oren et al. data, we calculated fold change values relative to all other time points for each gene using the Seurat FoldChange() function with cell identities being set to the time points. With these ranked gene lists, at each timepoint, we calculated the enrichment of the 72 gene resistance signature at all time points using GSEA from the fgsea R package^50^. We then plotted the normalized enrichment score for the four time points as a bar plot.

### Resistance signature functional enrichment

To identify which relevant pathways are enriched in the drug resistance signature, we utilized the clusterProfiler R package. The pathways we investigated for enrichment included mSigDB hallmarks, intratumoral heterogeneity meta-programs (MPs), and GO biological process terms. We performed a hypergeometric test via the enricher() function for hallmarks and MPs and the enrichGO() function for GO terms. The number of shared genes between a functional pathway and the geneset of interest is reported as counts.

### AUCell Threshold

For all applications of AUCell, to estimate a threshold for the raw AUCell score to designate a cell ‘active’ or ‘inactive’ for the gene signature, we created a background distribution of AUCell scores based on 100 control geneset with similar gene expression levels as the given geneset. To generate these control gene sets, we follow a procedure similar to that outlined in the AddModuleScore function in Seurat. We first assigned all expressed genes into ten bins based on their mean normalized expression and assigned each signature gene to a bin based on its normalized expression level. To generate a single control gene set, we picked a single gene at random from the same expression bin corresponding to each signature gene. The activity of the control gene set was then scored using AUCell, where the 95th percentile of the AUCell scores was stored as a putative activity threshold. This entire process was repeated 100 times, where the largest threshold computed is the final activity threshold. Any cell with an AUCell score higher than this threshold is considered to have an active resistance signature.

### Odds ratio for resistance signature activity in a cell cluster calculation

The odds ratio for each cluster is calculated as (# of active cells in cluster X/# of inactive cells in cluster X)/ (# of active cells not in cluster X/# of inactive cells not in cluster X) We use an odds ratio threshold of 1 and adjusted p-value threshold of 0.05 as the cutoff for resistance-activated clusters.

### Cancer hallmark heatmap creation

We scored every cell for each cancer hallmark geneset^17^ with AUCell then plotted the mean cluster score for each hallmark and the Z-scored ((0,1)-normalization) values for each hallmark across clusters within a cell line.

### Global RAC gene signatures

To acquire the gene signatures that define the RAC cells in each cell line we used the Seurat FindAllMarkers function to compare all RAC cells to all non-RAC cells. We further filtered the resulting differentially expressed genes for adjusted p value < 0.05 and average log2FC > 0. We sorted the genes by descending fold change and selected the top 200 genes.

### Global RAC signatures functional enrichment

To identify which relevant pathways are enriched in the global RAC signatures, we utilized the clusterProfiler R package. The pathways we investigated for enrichment included mSigDB hallmarks, intratumoral heterogeneity meta-programs (MPs), and GO biological process terms. We performed a hypergeometric test via the enricher() function for hallmarks and MPs and the enrichGO() function for GO terms. The number of shared genes between a functional pathway and the geneset of interest is reported as counts.

### Grouping the RACs across cell lines into superclusters

We first created a gene list that consists of the intersection of the top 2000 most variable genes in each cell line. With the resulting list of 3084 genes, for each gene, we calculated its differential mean cluster expression in a RAC relative to all other cells in each respective cell line. We then performed an all-by-all (for all clusters across the three cell lines) Spearman correlation of these differential gene expression vectors and hierarchically grouped the cell line-specific clusters. We defined superclusters as highly correlated groups of RACs including components from all three cell lines.

### Supercluster Consensus Signature creation

To create the signature to represent the supercluster spanning all three cell lines, we repeated the same process as the drug resistance signature using differentially expressed genes and RRA. We first selected differentially expressed genes (average log2FC > 0 and adjusted p value < 0.05) in each supercluster’s component RACs within their respective cell lines then sorted all gene lists by log2FC. Finally, we formed a single ranked list using Robust Rank Aggregation (RRA) and further filtered to retain all genes with an adjusted rank p-value < 0.05. For downregulated genes, the same process is repeated with a log2FC threshold of < 0.

### Supercluster Consensus Signature Functional enrichment

To identify which relevant pathways are enriched in the supercluster consensus signatures, we utilized the clusterProfiler R package. The pathways we investigated for enrichment included mSigDB hallmarks, intratumoral heterogeneity meta-programs (MPs), and GO biological process terms. We performed a hypergeometric test via the enricher() function for hallmarks and MPs and the enrichGO() function for GO terms. The number of shared genes between a functional pathway and the geneset of interest is reported as counts.

### Supercluster Cell Cycle phase plots

To calculate the percentage of cells in a supercluster that are in each of the cell cycle phases, we first scored all cells in each supercluster for gene signatures associated with each phase using the Seurat CellCycleScoring^51^ function. This function was modified to include an additional G0 signature derived from Wiecek et al.^52^. Each cell was assigned a phase based on the highest scoring cell cycle phase signature in that cell. With each cell assigned to a phase, we plotted the percentage of each phase for each supercluster.

### Drug class specific superclusters

To determine if cells treated with a specific drug class are overrepresented in the superclusters, we computed an odds ratio comparing the proportion of cells treated with a given drug class in the component clusters of each supercluster to the proportion of cells treated with the same drug but not in the supercluster component clusters within the respective cell lines.

### CRISPR Screening

A genome-wide CRISPR Brunello knockout (KO) lentiviral pooled library was used to assess genes required for resistance to the CHEK1/2 inhibitor, Prexasertib. The Brunello library contains 76,441 gRNAs targeting 19,114 genes and 1,000 unique non-targeting sgRNA controls (Addgene # 73178). First, OVCAR8 ovarian cancer cells were selected for stable Cas9 expression (Addgene # 52962-LV). Then, the Brunello sgRNA library was transduced at ∼0.3 MOI to ensure unique sgRNA distribution, and cells were selected with 5 μg/ml of puromycin for three days. The initial pooled cell population was split into separate study arms with an estimated 500x library coverage per condition. The treatment (30ng Prexasertib) and control (DMSO) arms were maintained in culture for 10 generations at a density required for library representation. Genomic DNA was extracted using the QIAmp DNA blood Cell Maxi Kit (Qiagen) according to the manufacturer’s protocol. The sgRNA barcodes were amplified using Taq polymerase and adapted for sequencing. The desired DNA product was purified with 6% TBE gel (Invitrogen) and samples were sequenced on an Illumina HiSeq2000. The sgRNA read count and hit calling were analyzed using MAGeCK (v0.5.7)^53^. Gene scoring and ranking were determined by Robust Rank Aggregation (RRA) with normalization relative to the distribution of non-targeting gRNAs. Genes that were negatively selected in the treatment arm compared to control were selected for downstream analysis.

### Kaplan-Meier plots

To create the Kaplan-Meier plots in this work we utilized the survminer R package. Samples were split into high and low expression groups based on top and bottom 50th percentiles.

### Cox regression

To model the effect of our resistance signatures on overall survival in TCGA data we performed Cox regression using the survival R package. For each sample we modeled overall survival with predictor variables being ssGSEA score for the current resistance signature, age, sex, and tumor purity (if available). We then plotted the hazard ratios for each cancer type as a forest plot.

### Scoring recurrence, pre-malignancy, and drug response data

For all three analyses involving scoring patient samples with global RAC or supercluster signatures, we utilized GSVA R package^29^. We scored each sample using the ssGSEA method and plotted the distribution of score between the two relevant groups of comparison.

### Creating resistance ortholog signatures

To create the drug resistance signature for *Candida auris*, we first obtained differential gene expression results between drug resistant and drug sensitive population from the original publication^13^. We then filtered for genes that are upregulated in the antifungal resistance population (log2FC > 0 and adjusted p-value < 0.05). For the drug resistance signature for *Escherichia coli,* we obtained differential gene expression results between treated and naive populations across 9 experiment using various unique antimicrobial drugs^12^. We next filtered for genes that are upregulated in each respective experiment (log2FC > 0 and adjusted p-value < 0.05). Finally, we sorted the 9 gene lists by log2FC and obtained a consensus signature by selecting genes that are upregulated in at least 5 of the 9 lists. Once both the resistance signatures were constructed for both *C. auris* and *E. coli*, we converted each gene to its orthologous match in the human genome using OMA Browser^34^ conversion tables.

### Scoring for resistance ortholog signature in superclusters

For both the *C. auris* and *E. coli* resistance ortholog signatures, we scored all of the cells in our early response data using AUCell. We then plotted the distribution of scores across all supercluster component clusters and non-RACs. We then compared the distributions for significance differences using Wilcoxon rank sum test.

### Functional Enrichment of conserved genes

To calculate the functional enrichment of the shared genes. Between human and *E. coli* or *C. auris*, we first identified all overlapping genes between the *E. coli* or *C. auris* resistance signature and the signature of particular superclusters that showed AUCell score enrichment for a given *E. coli* or *C. auris* resistance signature. With these lists of overlapping genes, we completed an overrepresentation analysis for hallmarks, ITH MPs, and GO processes as detailed above. The number of shared genes between a functional pathway and the geneset of interest is reported as counts.

## QUANTIFICATION AND STATISTICAL ANALYSIS

Error bars were calculated using fgsea function. Fisher tests were used for all odds ratio calculations. All two-group comparisons were conducted using Wilcoxon signed-rank test. Cox regression analyses used signature scores controlling for age, sex, and tumor purity.

