## Supplementary material for "Resistance Signatures Manifested in Early Drug Response in Cancer and Across Species": Document S1

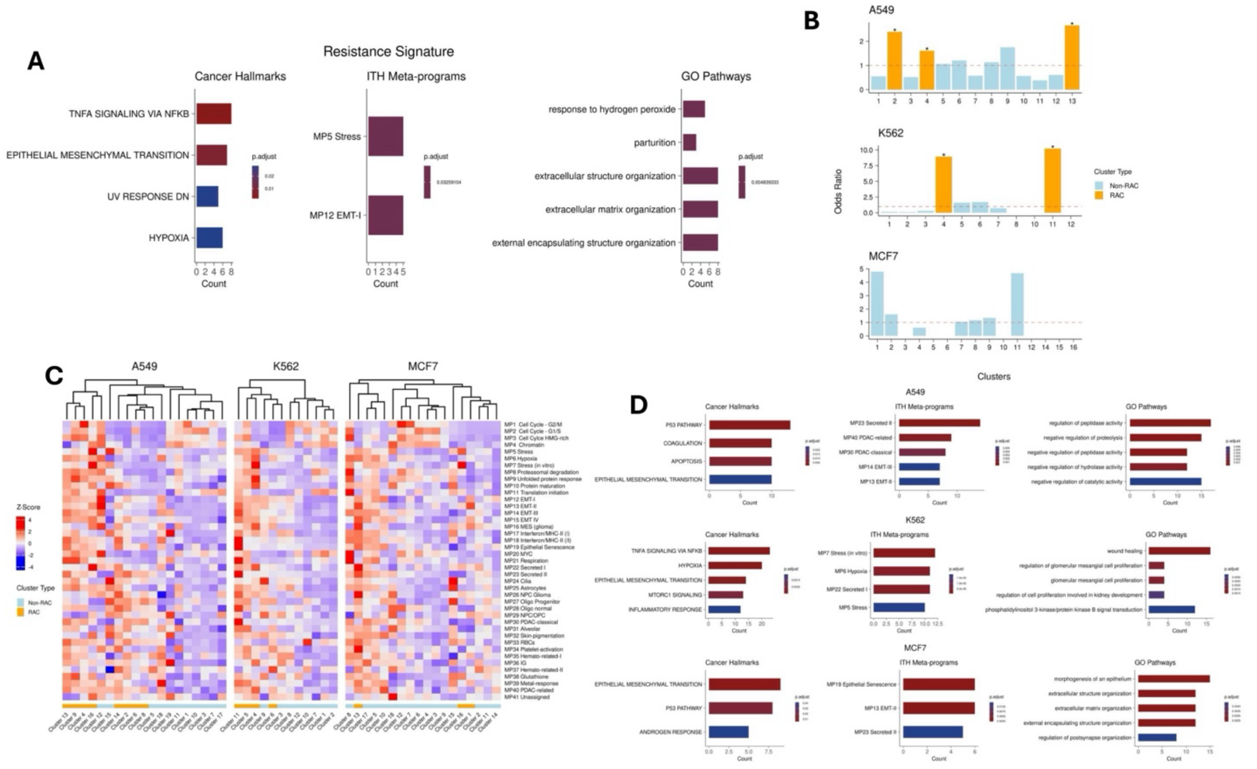


### Figure S1. Characterization of Resistance

(A) Pathway enrichment of drug resistance signature. Each plot shows the overrepresentation of cancer hallmarks, ITH meta-programs, and GO pathways respectively. (B) Pre-treatment resistant active to inactive cell odds ratio of each cluster in each cell line. The y axis represents the odds ratio value and each bar represents a transcriptional cluster. Any cluster with an odds ratio greater than 1.5 and a p-value less than 0.05 is considered a Resistance Activated Cluster (RAC) and is colored orange, while the remaining clusters are non-RACs and are colored blue. (C) Intratumor Heterogeneity Meta-program mean AUCell score for all clusters in each cell line. The rows represent the different ITH MP genesets and the columns represent each transcriptional cluster, separately for each cell line. The colors in the bottom row signifies which cluster is a Resistance Activated Cluster (RAC) (orange = RAC, blue = Non-RAC). (D) Enrichment of the global RAC signature in each cell line. The columns from left to right represent cancer hallmarks, ITH meta-programs, and GO pathways respectively. Within each column, the rows represent the three cell lines. Count values indicate the number of shared genes between the global RAC signature and the enriched pathway.

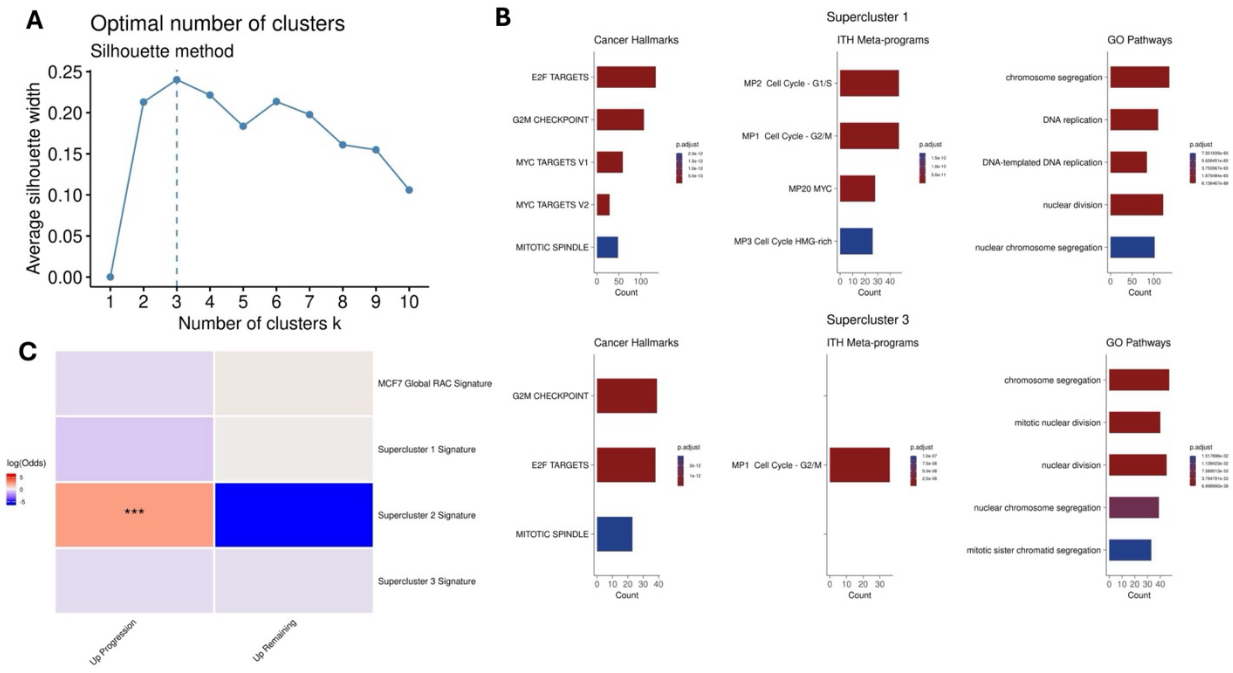


### Figure S2. Supercluster Enrichment

(A) Optimal number of superclusters. Line plot showing the silhouette score for each number of superclusters. Vertical line drawn at maximum value indicating the optimal number of superclusters. (B) Supercluster Downregulated Signatures Enrichment. The columns from left to right represent cancer hallmarks, ITH meta-programs, and GO pathways respectively. Within each column, the rows represent the three superclusters. Count values indicate the number of shared genes between the supercluster downregulated signature and the enriched pathway. (C) Shared progression-associated genes. Displays the log odds ratio of overrepresentation test observing the overlapping genes between the four resistance signatures and progression-associated genes.
